# High-resolution imaging and manipulation of endogenous AMPA receptor surface mobility during synaptic plasticity and learning

**DOI:** 10.1101/2021.09.21.461215

**Authors:** Angela M. Getz, Mathieu Ducros, Christelle Breillat, Aurélie Lampin-Saint-Amaux, Sophie Daburon, Urielle François, Agata Nowacka, Mónica Fernández-Monreal, Eric Hosy, Frédéric Lanore, Hanna Zieger, Matthieu Sainlos, Yann Humeau, Daniel Choquet

**Author notes:** Corresponding Author: Daniel Choquet. These authors contributed equally to this work (Yann Humeau, Daniel Choquet).

## Abstract

Regulation of synaptic neurotransmitter receptor content is a fundamental mechanism for tuning synaptic efficacy during experience-dependent plasticity and behavioral adaptation. However, experimental approaches to track and modify receptor movements in integrated experimental systems are limited. Exploiting AMPA-type glutamate receptors (AMPAR) as a model, we generated a knock-in mouse expressing the biotin acceptor peptide (AP) tag on the GluA2 extracellular N-terminus. Cell-specific introduction of biotin ligase allows the use of monovalent or tetravalent avidin variants to respectively monitor or manipulate the surface mobility of endogenous AMPAR containing biotinylated AP-GluA2 in neuronal subsets. AMPAR immobilization precluded the expression of long-term potentiation and formation of contextual fear memory, allowing for target-specific control of the expression of synaptic plasticity and animal behavior. The AP tag knock-in model offers unprecedented access to resolve and control the spatiotemporal dynamics of endogenous receptors, and opens new avenues to study the molecular mechanisms of synaptic plasticity and learning.

## INTRODUCTION

Changes in synaptic transmission efficacy and network connectivity, particularly in the context of experience-dependent synaptic plasticity, are at the core of behavioral adaptation, learning and memory (Humeau and Choquet, 2019; Josselyn and Tonegawa, 2020). This has brought center stage the need to understand the cellular and molecular mechanisms that modulate synaptic function. Among the many pre- and post-synaptic mechanisms that control synaptic gain, regulation of the number, type, and nanoscale organization of neurotransmitter receptors at the postsynaptic density (PSD) has emerged as a principle determinant of the amplitude of both excitatory and inhibitory synaptic responses (Choquet and Triller, 2013; Groc and Choquet, 2020). This is established by a complex interplay between a series of trafficking processes, including intracellular transport, exo- and endocytosis, lateral diffusion on the cell surface, and reversible stabilization at the PSD through interactions with scaffolding elements. Altogether, this results in a dynamic equilibrium of receptors distributed amongst different subcellular compartments that ultimately sets their number in front of transmitter release sites.

Molecular replacement strategies or overexpression approaches introducing modified receptor subunits (e.g. chimeras, truncations, super-ecliptic pHluorin (SEP) tags) have been widely used to study the contributions of receptor trafficking mechanisms to synaptic plasticity (Chen et al., 2021; Granger et al., 2013; Kellermayer et al., 2018; Makino and Malinow, 2009; Petrini et al., 2014; Zhang et al., 2015). Notwithstanding their extensive use, these approaches face a number of experimental limitations, including disrupting the set point of the ‘diffusion-trapping’ equilibrium through changes in receptor subunit composition, synaptic targeting, alterations of the receptor surface pool, and saturation of synaptic anchoring sites. Such off-target effects are likely to produce measurement artefacts, while allowing only a handful of neurons in a circuit to be assayed at a given time. This makes them of limited use for addressing experimental questions at the level of integrated neuronal circuits under endogenous conditions, or with behavioral paradigms *in vivo*.

An important feature of the control of receptor numbers at synapses is that sites of receptor endo- and exocytosis are primarily extrasynaptic (Blanpied et al., 2002; Lin et al., 2009; Petrini et al., 2009). Consequently, the primary pathway for the addition or removal of synaptic receptors is through surface movement powered by Brownian diffusion (Ashby et al., 2006; Borgdorff and Choquet, 2002; Meier et al., 2001). Controlling receptor surface diffusion has emerged as a powerful avenue towards artificial regulation of synaptic plasticity by preventing variations in receptor numbers at synapses. The use of antibodies against extracellular domains of endogenous receptor subunits (e.g. GABA_A_, NMDA, AMPA, kainate receptors) has proven an effective method to control their movements on the neuronal surface through the crosslinking effects introduced by the divalent binding domains of antibodies (de Luca et al., 2017; Dupuis et al., 2014; Heine et al., 2008; Penn et al., 2017; Polenghi et al., 2020). Importantly, extracellular crosslinkers provide a specific tool to manipulate receptor mobility without altering basal synaptic transmission or circuit function. Such approaches have established the pivotal role of AMPA receptor (AMPAR) surface trafficking in short-term and canonical long-term plasticity (LTP), as well as the link between variations in synaptic receptor numbers and higher brain functions such as cued fear learning and whisker-dependent somatosensory behavior (Campelo et al., 2020; Heine et al., 2008; Penn et al., 2017; Potier et al., 2016). The ability to visualize and control receptor fluxes during plasticity could potentially allow to study the precise timing and contributions of changes in synaptic receptor content during various higher brain functions such as working memory or memory engrams formation. However, progress in this area has been hindered by the lack of appropriate experimental tools that can specifically monitor or manipulate receptor dynamics at the level of single synaptic contacts or in particular subclasses of neurons within a circuit.

There is therefore a great need to develop new molecular tools that will allow to overcome the aforementioned experimental limitations. An ideal approach would be one that allows for target-specific monitoring and manipulation of endogenous receptor surface mobility dynamics, with resolution that can be tuned to the level of single molecules, individual synapses, or integrated synaptic networks, while preserving endogenous receptor composition and function. The avidin-biotin system for labeling cell surface proteins has recently emerged as a promising candidate for such an approach (Howarth et al., 2005; Howarth and Ting, 2008). Proteins tagged with the 15 amino acid biotin acceptor peptide (AP) sequence are selectively biotinylated in the endoplasmic reticulum (ER) upon the expression of biotin ligase (BirA) from *E. coli* modified to contain an ER retention sequence (BirA^ER^), or on the cell surface upon the addition of soluble recombinant BirA (sBirA). Biotinylated proteins on the cell surface are recognized by the avidin family of small, high-affinity biotin binding proteins, including StreptAvidin and NeutrAvidin (SA/NA; ∼6 nm), or monomeric StreptAvidin (mSA; ∼3 nm) (Chamma et al., 2017).

A number of features make the avidin-biotin system particularly well suited for studying the surface mobility dynamics and nanoscale organization of synaptic proteins in integrated systems. First, distinct molecular strategies to introduce BirA allow time-controlled and target-specific AP tag biotinylation and avidin recognition in a constitutive knock-in (KI) model where all neurons express the AP-tagged protein. Second, addition of the small AP tag sequence is unlikely to have a substantial impact on protein expression, structure, function, or trafficking. Third, the small size and high affinity of biotin binding proteins offer enhanced synaptic access, labeling stability, and signal intensity over antibody- or fluorescent protein conjugate-based approaches. Fourth, thanks to the availability of monovalent (mSA) and tetravalent (NA, SA) variants of biotin binding proteins, AP tagging presents a molecular strategy that can be used either to monitor the surface mobility dynamics and nanoscale organization of individual proteins, or to efficiently manipulate surface diffusion dynamics by crosslinking-dependent immobilization (Chamma et al., 2016; Penn et al., 2017).

Here, we report the development and characterization of an AP-GluA2 KI mouse model where *Gria2* was modified by CRISPR-Cas9 genome editing to introduce the AP tag to the extracellular N terminus of the GluA2 AMPAR subunit. The expression, localization and function of GluA2 was not affected by the modification, and the synaptic physiology and behavioral phenotype of KI animals was indistinguishable from wild-type (WT). We developed in parallel a molecular toolkit that allows the biotinylation of AP-GluA2 to be tailored to a variety of spatiotemporal resolutions and experimental preparations, ranging from super-resolution imaging of single molecules to manipulating integrated circuits in behaving animals. Development of a custom lattice light sheet microscope (LLSM) with a photostimulation module (PSM) enabled us to perform fluorescence recovery after photobleaching (FRAP) imaging of biotinylated AP-GluA2 with monovalent or tetravalent avidin probes to measure the native surface mobility and crosslinking-dependent immobilization of endogenous AMPAR in brain slices. In acute slices, NA crosslinking of biotinylated AP-GluA2 (bAP-GluA2)-containing AMPAR blocked the expression of LTP at Schaffer collaterals, and delivery of NA into the CA1 region *in vivo* blocked the formation of contextual fear memory. This experimental model affords unparalleled resolution and control over the spatiotemporal dynamics of endogenous AMPAR for integrated physiological studies.

## RESULTS

### Target-specific detection of surface biotinylated AP-GluA2 in organized brain tissue

We used CRISPR-Cas9 genome editing of *Gria2* to introduce the biotin acceptor peptide sequence (AP; GLNDIFEAQKIEWHE) and a downstream tobacco etch virus protease consensus sequence (TEV; ENLYFQG) onto the GluA2 N-terminus (Figure 1A, supplementary Figure S1). The AP tag enables endogenous GluA2-containing AMPAR to be labeled with avidin probes upon enzymatic biotinylation by BirA, while the TEV site allows the KI sequence to be enzymatically removed for reversible surface labeling and crosslinking. To achieve target-specific biotinylation of AP-GluA2, we developed a variety of molecular approaches via the chronic expression of ER-resident BirA (BirA^ER^) by adeno-associated virus (AAV) transduction or single cell electroporation (SCE), or the acute application of soluble recombinant BirA (sBirA) (Figure 1A). For applications requiring sparse labeling, we developed a two component AAV viral system, with one virus encoding endoplasmic BirA^ER^ with Cre recombinase following an IRES site (BirA^ER^-Cre), used at a low concentration with a second virus encoding a floxed eGFP reporter. We also used an alternative plasmid system encoding BirA^ER^ IRES eGFP (BirA^ER^-eGFP) (see Methods). The design of these constructs allows the reporter to be easily exchanged, thereby facilitating coupling the avidin-biotin system to various functional assays with fluorescent reporters. We used AAV microinjection or SCE of CA1 pyramidal neurons in organotypic hippocampal slices cultured with 10 µM biotin supplementation, and performed live-labeling with dye-conjugated NA to detect surface biotinylated AP-GluA2 (bAP-GluA2). For controls, we mirrored the above conditions in WT slices, or in AP-GluA2 KI slices without BirA^ER^, using an AAV encoding Cre alone or a plasmid encoding IRES eGFP. NA labeling was specific to bAP-GluA2 in KI slices with BirA^ER^, and well correlated with the eGFP reporter (Figure 1B-G). TEV incubation efficiently removed the extracellular NA label within 10 min (Figure 1B-D, Figure S2). To define the experimental timeframe for this system we monitored the expression of bAP-GluA2 over time in culture, and observed saturation after 12 days *in vitro* (DIV) with AAV-mediated expression of BirA^ER^ (Figure 1E, Figure S3), and after 6 DIV with plasmid-mediated expression of BirA^ER^ (Figure 1G, Figure S4).

**Figure 1:**
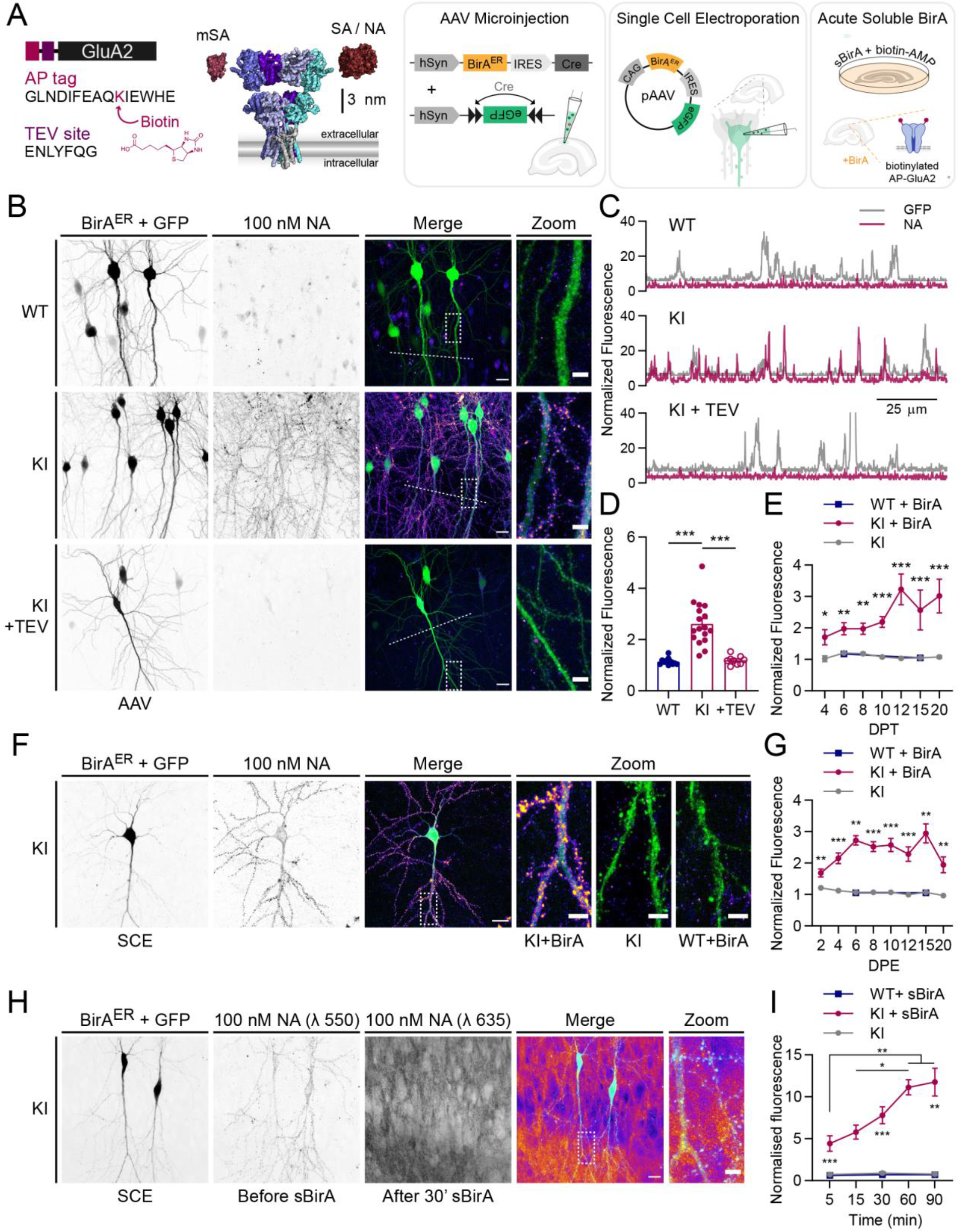
Target-specific surface labelling of AMPAR by biotin binding proteins in an AP-GluA2 KI model. (**A**). Schematic representation of the AP-GluA2 KI modified extracellular N-terminus, containing an AP tag and TEV protease consensus sequence, with a scaled model of AMPAR with monovalent (mSA) or tetravalent (SA or NA) biotin binding proteins (left), and the three molecular strategies to biotinylate AP-GluA2: long-term expression of BirA^ER^ by AAV transduction; plasmid electroporation; or acute incubation with sBirA (right). (**B**). Representative confocal images of CA1 pyramidal neurons in WT or AP-GluA2 KI slice cultures transduced with BirA^ER^-Cre + FLEx eGFP AAV, incubated with NA-DyLight 633 to label surface localized bAP-GluA2 (top, middle), then TEV protease to cleave the AP tag and remove the NA surface label (bottom). (**C**). Linescans (dashed lines in b) reveal extent of NA colocalization with the eGFP reporter (Pearson correlation; WT: R=0.0010, *P*=0.9686; KI: R=0.3359, \*\*\**P*<0.0001; KI+TEV: R=0.0371, *P*=0.2195). (**D**). Normalized fluorescence intensity of NA-DyLight 633, coincident with the eGFP reporter. N≥9. \*\*\**P*≤0.0001 (Welch’s ANOVA, F=23.99, *P*<0.0001; Dunnett’s post-hoc test). (**E**). Time course of bAP-GluA2 expression in KI slices transduced with BirA^ER^-Cre + FLEx eGFP AAV at DIV1, relative to WT (+BirA) or Cre alone (-BirA) control. DPT, days post transduction. Normalized fluorescence intensity of NA-DyLight 633, coincident with the eGFP reporter. N≥3. *-\*\*\**P*≤0.0201 (Mann-Whitney U-test, unpaired T-test or Kruskal-Wallis tests; F≥12.44, *P*≤0.002, Dunn’s post-hoc test). (**F**). CA1 pyramidal neurons in AP-GluA2 KI or WT slices electroporated with BirA^ER^-eGFP or eGFP alone control plasmids at DIV3. (**G**). Time course of bAP-GluA2 expression in KI slices electroporated with BirA^ER^-eGFP, relative to WT (+BirA) or eGFP alone (-BirA) control. DPE, days post electroporation. N≥5. **-\*\*\**P*≤0.0098 (Mann-Whitney U-test, unpaired T-test, Kruskal-Wallis test or Welch’s ANOVA; F≥16.22, *P*≤0.0006, Dunn’s or Dunnett’s post-hoc test). (**H**). Two-color labeling of BirA^ER^-eGFP electroporated KI slices, first with NA-DyLight 550, then with NA-STAR 635P after 30’ incubation with sBirA, which rapidly biotinylates surface AP-GluA2. (**I**). Time course of bAP-GluA2 expression in sBirA incubated KI slices, relative to WT or biotin-AMP alone control. N≥5. *-\*\*\**P*≤0.042 (Kruskal-Wallis test or Welch’s ANOVA; F≥14.69, *P*≤0.0003, Dunn’s or Dunnett’s post-hoc test). Scale bars, 20 and 5 µm. Error bars, SEM. See also Supplementary Figures S1-S6.

For bulk labeling applications, we developed an acute approach using incubation with sBirA and biotin-AMP (adenosine 5′-monophosphate), where detection of bAP-GluA2 saturated after 60 min (Figure 1H-I, Figure S6), and a chronic approach with an AAV encoding BirA^ER^ IRES eGFP (BirA^ER^-eGFP), where essentially all neurons within the infection zone expressed bAP-GluA2 (Figure S5 and Figure 6B, 7B). It is of interest to note that this AP tag KI system is amenable to the characterization of distinct populations of surface AMPAR using two-color avidin labeling before and after an experimental stimulus. Here, for example, we used differentially dye-coupled NA to identify surface bAP-GluA2 before and after acute biotinylation by sBirA (Figure 1H). The long-term BirA^ER^ gene delivery strategies and the rapid acute sBirA approaches therefore offer access to two distinct temporal windows, making the AP-GluA2 KI model amenable to a variety of experimental approaches. Taken together, these results demonstrate the high degree of specificity and flexibility of this system for labeling endogenous AMPAR in organized brain tissue.

### Monitoring and manipulating endogenous AMPAR surface mobility

The small size of biotin binding proteins affords enhanced labeling accessibility for confined environments such as in organized tissue or at the synapse (Chamma et al., 2016). However, live high-resolution imaging of endogenous proteins dynamics in optically-scattering tissue preparations represents a considerable challenge. To circumvent photobleaching, phototoxicity, and optical scattering constraints, we developed a lattice light sheet microscope (LLSM) with a photostimulation module (PSM) to determine the mobility of bAP-GluA2 by measuring the rate of fluorescence recovery after photobleaching (FRAP) (Figure S7). In BirA^ER^-eGFP or BirA^ER^-Cre + eGFP transduced KI organotypic slices, we labeled surface bAP-GluA2 with 400 nM mSA or 100 nM NA conjugated to the fluorophore STAR 635P (Figure 2A). Regions of interest (ROIs) on CA1 pyramidal neuron spines and dendrites in the *stratum radiatum* were photobleached and fluorescence recovery was followed for ∼250 s to measure the mobility of synaptic and extrasynaptic AMPAR. Using mSA, we found that the fraction of mobile AMPAR was 0.27 ± 0.03 at spines, and 0.50 ± 0.05 on dendrites. With NA, the mobile fraction was reduced to 0.01 ± 0.03 at spines, and 0.22 ± 0.03 on dendrites (Figure 2B-F). This indicates that (i) mSA labeling of bAP-GluA2 affords access to monitoring the diffusion dynamics of endogenous AMPAR in organized brain tissue, and (ii) NA-mediated crosslinking of bAP-GluA2 efficiently reduces the surface diffusion of endogenous AMPAR and eliminates their exchange between synaptic and extrasynaptic sites.

**Figure 2:**
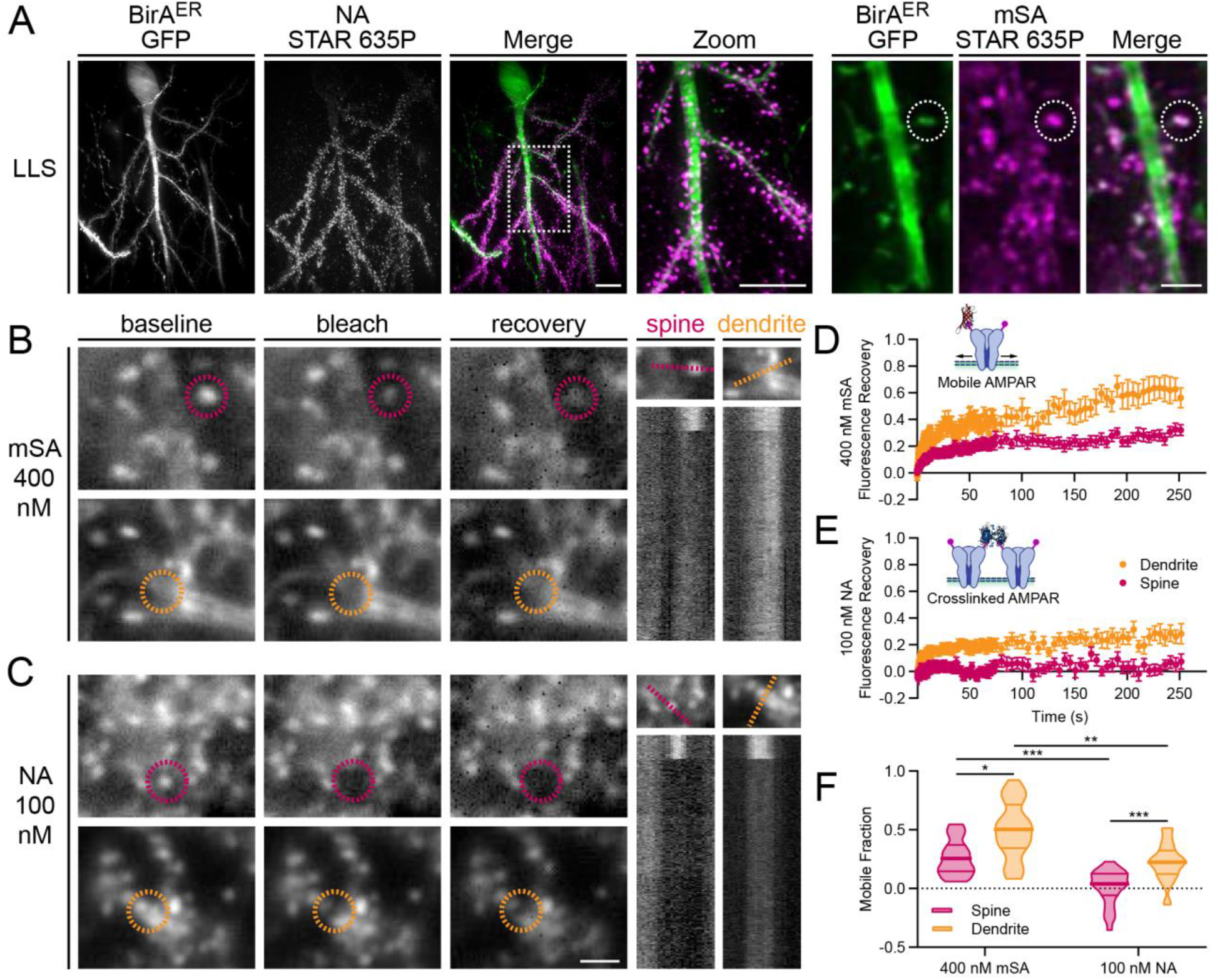
LLSM-FRAP with biotin binding proteins reveals surface mobility and immobilization of AP-GluA2-containing AMPAR. (**A**). Representative 3D reconstructed LLSM images of CA1 pyramidal neurons in AP-GluA2 KI slice cultures transduced with BirA^ER^-Cre + FLEx eGFP or BirA^ER^-eGFP AAV, incubated with NA (left) or mSA (right) to label surface localized bAP-GluA2. Scale bars, 10 and 2 µm. (**B-C**). FRAP experiments performed on a custom LLSM with PSM in AP-GluA2 KI slices labeled with mSA-STAR 635P (**B**) or NA-STAR 635P (**C**). Representative images show spine or dendrite ROIs (dashed circles) before (baseline; −1s) and after targeted photobleaching (bleach; +1s), and diffusion-dependent recovery (recovery; +70s). Scale bar, 2 µm. Kymographs illustrate fluorescence recovery profiles of the targeted ROIs during the acquisition (dashed line; ∼250s). (**D-E**). Normalized mean fluorescence recovery curves for bAP-GluA2 labeled with monovalent mSA reveals synaptic and extrasynaptic mobility of endogenous AMPAR (**D**), or tetravalent NA, which immobilizes surface AMPAR by crosslinking of bAP-GluA2 (**E**). N≥25. 52-88% of FRAP ROIs were maintained in the LLS excitation plane and followed for the full length of the acquisition. Error bars, SEM. (**F**). Quantification of the AMPAR mobile fraction by curve-fitting of individual FRAP recovery profiles, plot shows median with first and third quartiles. N≥25. *-\*\*\**P*≤0.0333 (Kruskal-Wallis test; F=53.60, *P*<0.0001; Dunn’s post-hoc test). See also Supplementary Figures S7-S9.

As the crosslinking effects of certain autoantibodies against GluA2 have been reported to trigger the internalization of surface bound AMPAR (Haselmann et al., 2018; Peng et al., 2015), we next sought to characterize the impact of NA-mediated crosslinking on AMPAR internalization. To this end, we used kymograph plot profiles to analyze the number of internalized vesicles observed passing through a ∼ 5 µm dendritic segment during LLSM FRAP acquisitions, and found no significant difference between mSA and NA labeled bAP-GluA2 (Figure S8). We then used tetrodotoxin (TTX) or NMDA treatment on mSA or NA labeled slices to respectively decrease or increase the rate of activity-dependent AMPAR internalization. After 30 min we used TEV to cleave surface labeled bAP-GluA2 to reveal the fraction of internalized AMPAR. Again, we found no significant difference between mSA and NA labeled bAP-GluA2 (Figure S9). This suggests that NA is an efficient molecular tool to control AMPAR surface diffusion with minimal impact on receptor surface localization and recycling dynamics.

### High-resolution imaging of AP-GluA2 synaptic organization and surface mobility

As the N-terminal domain of AMPAR subunits has been reported to influence synaptic organization and surface mobility dynamics (Diaz-Alonso et al., 2017; Watson et al., 2017), we next performed direct stochastic optical reconstruction microscopy (dSTORM) and universal point accumulation for imaging in nanoscale topography (uPAINT) super-resolution imaging to characterize AMPAR synaptic nanoscale organization and surface diffusion in dissociated hippocampal neuron cultures from AP-GluA2 KI or WT mice. Surface AMPAR were live-labeled with the 15F1 GluA2 antibody before fixation and observed by dSTORM using an Alexa 647 coupled secondary antibody. The number of AMPAR were estimated based on the blinking properties of the fluorophore (Nair et al., 2013). We found no difference in the number of AMPAR at spines or synaptic nanoclusters between WT and KI cultures (Figure 3A-C). For uPAINT, surface GluA2 were live-labeled with a low concentration of SeTau 647-coupled 15F1 to measure total surface AMPAR diffusion. We found no difference in the distribution of diffusion coefficients or the fraction of mobile receptors between KI and WT cultures (Figure 3D-F). This suggests that the modified N-terminal sequence in the AP-GluA2 KI model does not impact AMPAR synaptic organization and mobility.

**Figure 3:**
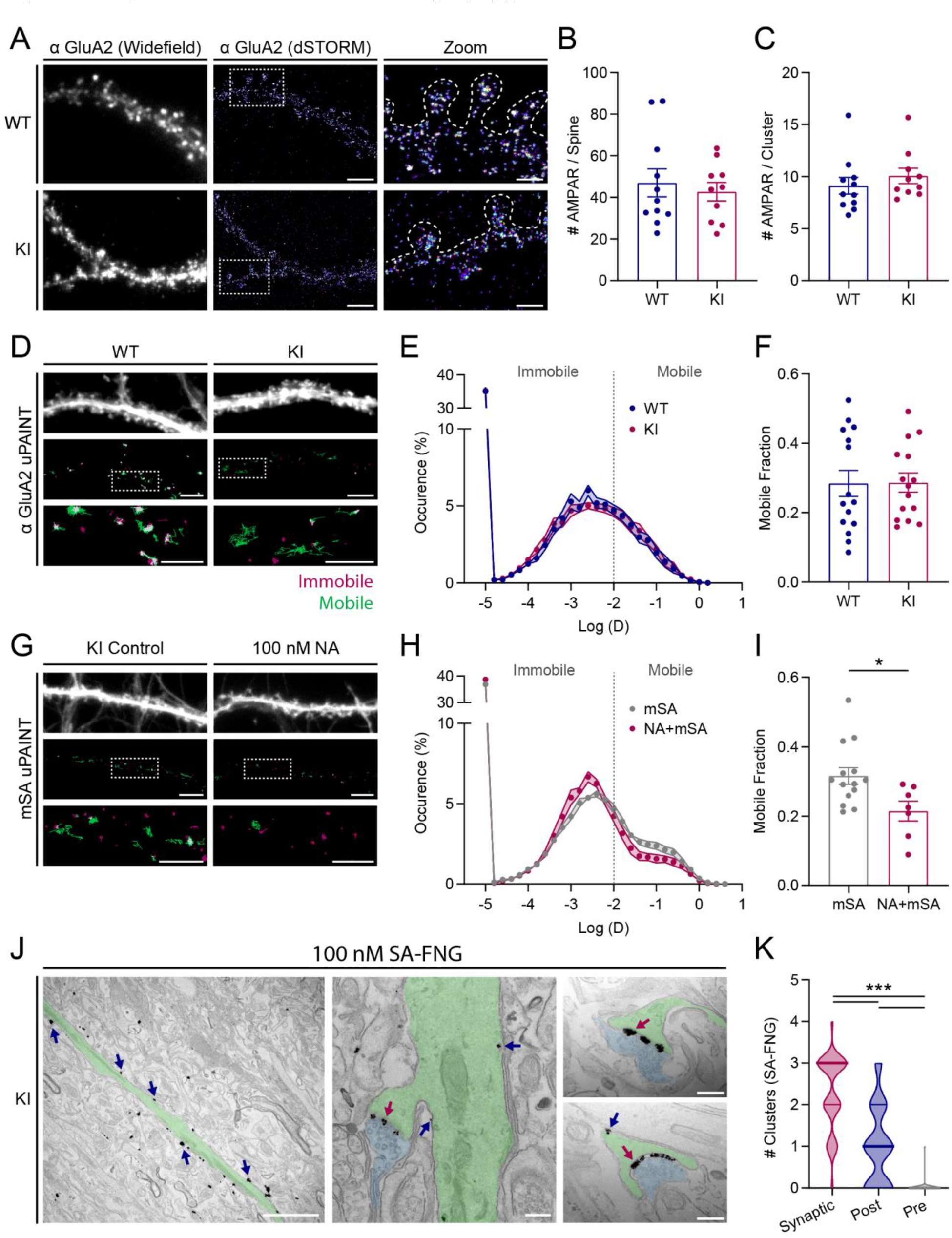
Super-resolution and TEM imaging approaches with AP-GluA2. (**A**). Representative widefield and super-resolved dSTORM images of WT and KI hippocampal neurons in primary dissociated culture, live-labeled with α-GluA2. (**B-C**). Number of AMPARs on individual spines (**B**) and in synaptic nanodomain clusters (**C**) estimated by counting the single emitters, plots show median per cell. N≥10. *P*≥0.1971 (Mann-Whitney U-tests). (**D**). Representative images of WT and KI hippocampal neurons in primary dissociated culture, transduced with eGFP AAV (top, widefield image) and live-labeled with α-GluA2-SeTau 647 (middle, reconstructed uPAINT trajectories; bottom, zoom of boxed region; mobile trajectories in green, immobile in red [Log(D) ≥ or ≤ −2, respectively]). (**E-F**). Average distribution of the logarithm of diffusion (**E**), and ratio of the mobile over immobile fraction (**F**). N=15. *P*=0.9675 (unpaired T-test). (**G**). Representative widefield and uPAINT images of KI hippocampal neurons in primary dissociated culture transduced with BirA^ER^-eGFP AAV, incubated with vehicle control or 100 nM unconjugated NA. A low concentration of mSA-STAR 635P (7.8 nM) was used for the recording of trajectories of bAP-GluA2. (**H-I**). Average distribution and ratio, as in (**E-F**). N≥7. \**P*=0.0193 (unpaired T-test). Scale bars, 5 and 2 µm. Error bars, SEM. (**J**). Representative TEM images of bAP-GluA2 in CA1 pyramidal neurons from AP-GluA2 KI slice cultures transduced with BirA^ER^-eGFP AAV. Slices were incubated with 100 nM SA conjugated to AlexaFluor 546 FluoroNanoGold (SA-FNG), then fixed and processed with silver enhancement to enlarge the gold particles for visualization. Left panel shows an apical dendrite (shaded green) with multiple SA-FNG clusters (arrows). Center panel shows a spine and dendrite with a presynaptic bouton (shaded blue), with multiple synaptic (red arrow) and extrasynaptic SA-FNG clusters (blue arrows). Right panel shows representative images used to quantify SA-FNG clusters, localized to the synaptic cleft (red arrow), postsynaptic spine (blue arrow), or presynaptic bouton (not shown/infrequently detected). Scale bars, 2 µm and 200 nm. (**K**). Subcellular distribution of SA-FNG clusters, plot shows median with first and third quartiles. N=39 synapses from three independent experiments. \*\*\**P*<0.0001 (Kruskal-Wallis test; F=74.88, *P*<0.0001; Dunn’s post-hoc test). See also Supplementary Figure S10.

We then used a low concentration of STAR 635P-coupled mSA to monitor AMPAR surface diffusion by tracking bAP-GluA2. To evaluate the impact of NA crosslinking on AMPAR lateral diffusion, we pretreated cultures with unlabeled NA before the application of mSA, reasoning that remaining free biotin sites within the pool of NA-crosslinked surface AMPAR would be accessible to dye-conjugated mSA. Indeed, NA application shifted the distribution of bAP-GluA2 diffusion coefficients towards lower values with a concomitant decrease in the fraction of mobile receptors (Figure 3G-I), consistent with our observations of NA-mediated immobilization in organotypic slices by LLSM-FRAP (see Figure 2E-F).

Next, we took advantage of the large catalogue of commercially available SA conjugates and used FluoroNanogold (SA-FNG) to characterize the distribution of bAP-GluA2-containing AMPAR in organotypic slices by transmission electron microscopy (TEM) (Figure S10). We quantified the distribution of silver-enhanced SA-FNG clusters on CA1 *stratum radiatum* synaptic micrographs, and found that clusters were predominantly localized to the synaptic cleft and postsynaptic spine membranes, but infrequently associated with presynaptic bouton membranes identified by synaptic vesicle content (Figure 3J-K). Taken together, these observations demonstrate the utility of the AP-GluA2 KI model and small avidin probes to study the nanoscale organization and diffusion dynamics of endogenous AMPAR with high-resolution imaging approaches.

### Biochemical characterization confirms wild-type expression levels of AP-GluA2

To characterize AP-GluA2 protein expression in the KI model, we performed Western blot analysis of KI or WT brain protein lysates. The assay of several synaptic proteins indicated no differences in expression, except for AP-GluA2, which surprisingly appeared as a smeared double band with reduced apparent expression when quantified (Figure 4A-B, Figure S11). However, when protein samples were incubated with TEV to cleave the AP tag, GluA2 resolved to a single band in the AP-GluA2 KI with the same relative expression as in WT (Figure 4C-D, Figure S11). As the observed shift in the apparent molecular weight of AP-GluA2 is larger than would be expected for the addition of 48 amino acids encoding the AP-TEV and linker sequence (∼5 kDa), we used various glycosidases to evaluate the glycosylation state AP-GluA2. We found that deglycosylation did not impact the double banding pattern (Figure S12A). While both upper and lower GluA2 bands were recognized by an AP tag antibody, the overlap between α-AP tag and α-GluA2 signals was incomplete, with the lower part of the faster migrating GluA2 band not recognized by the α-AP tag (Figure 4E-F, Figure S12B). When we incubated the protein samples with sBirA and biotin-AMP, SA binding to bAP-GluA2 overlapped with the GluA2 upper band and intermediate smear, but was absent from the lower band (Figure 4G-H,K, Figure S13). These observations suggest that the lower band represents an N-terminal degradation product of AP-GluA2 which has lost the biotinylated lysine. We then performed subcellular fractionation to assay the distribution of AP-GluA2 protein, and found that the upper band of AP-GluA2 was enriched in the synaptic membrane fraction, whereas the lower band was enriched in the vesicular fraction (Figure 4H-J,L, Figure S14). This indicates that most surface GluA2 in the KI model carry the AP tag conducive to biotinylation by BirA, and therefore recognition by biotin binding proteins.

**Figure 4:**
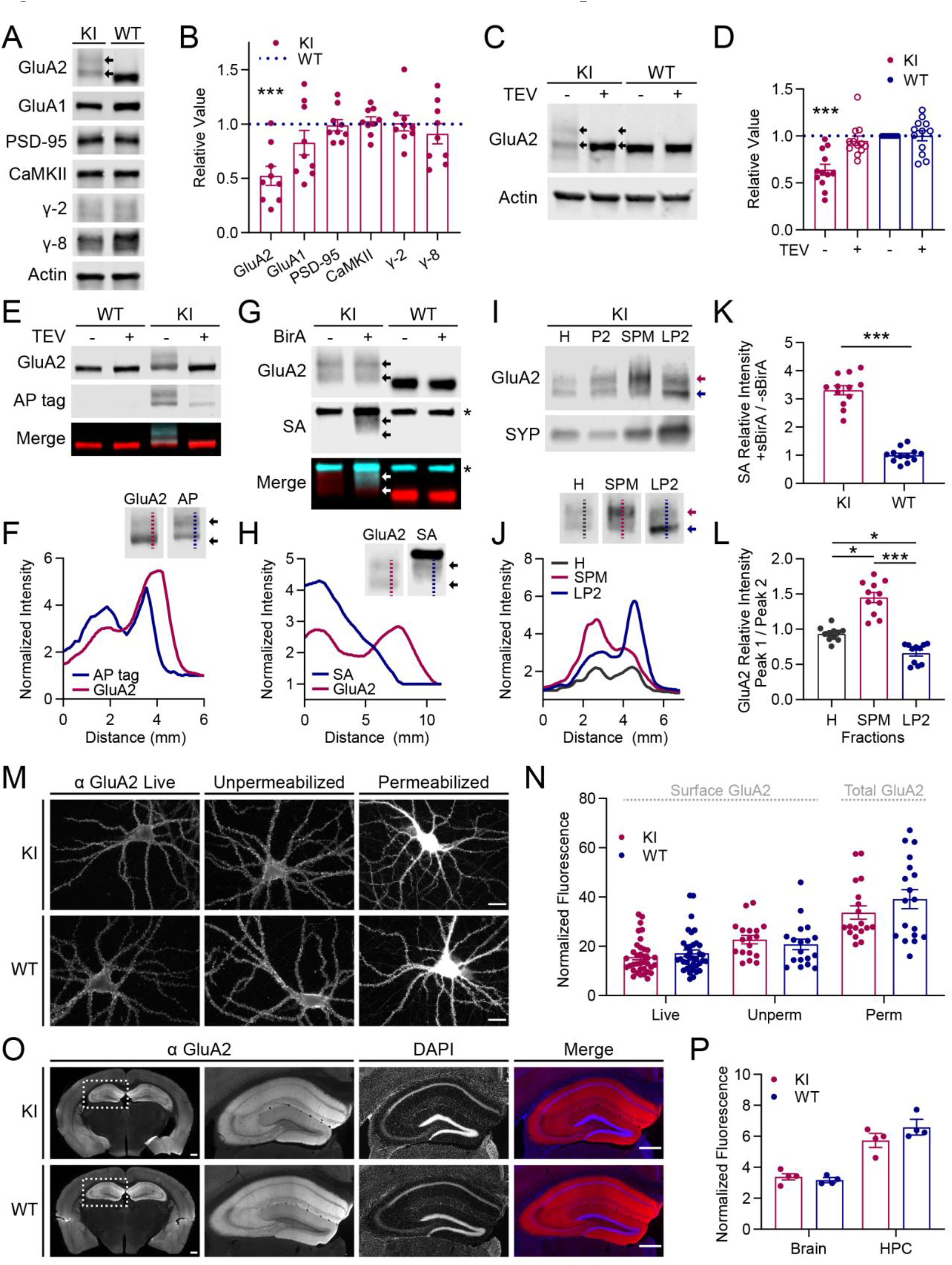
Biochemical characterization of AP-GluA2 expression and localization. (**A**). Representative Western blots of AP-GluA2 KI or WT protein samples (whole brain or hippocampal lysate). Double banding is observed for AP-GluA2 KI (arrows). (**B**). Quantification of KI protein expression relative to WT and normalized to β-actin loading control. N=9. \*\*\**P*=0.0006 (one sample T-tests). (**C**). *In vitro* incubation of KI or WT protein samples with TEV protease cleaves the AP tag and resolves GluA2 to a single band (arrows) with the same relative expression as WT. (**D**). Quantification of GluA2 expression relative to WT (-TEV) and normalized to β-actin loading control. N=12. \*\*\**P*=0.0005 (Wilcoxon Signed Rank test). (**E**). Dual labeling of AP tag and GluA2 in WT or KI protein samples with or without TEV protease. (**F**). Linescans (dashed lines in inserts) reveal partial colocalization of AP tag and lower GluA2 band (arrows). (**G**). *In vitro* incubation of KI or WT protein samples with sBirA + biotin-AMP, SA binds bAP-GluA2 in the upper GluA2 band (arrows). Asterisk denotes SA binding to endogenous biotin binding proteins. (**H**). Linescans reveal colocalization of SA with the upper but not lower GluA2 band. (**I**). Subcellular fractionation of hippocampal lysate (H, homogenate; P2, crude membranes) shows enrichment of the upper AP-GluA2 band in the synaptic plasma membrane fraction (SPM) and the lower band in the vesicular fraction (LP2). (**J**). Linescans show differential sorting of AP-GluA2 populations to the membrane (SPM, red line and arrow) and intracellular vesicles (LP2, blue line and arrow). (**K**). Quantification of SA binding (+ relative to - sBirA). N=12. \*\*\**P*<0.0001 (unpaired T-test). (**L**). Quantification of relative AP-GluA2 distribution (upper, peak 1; lower, peak 2) in linescans from blots of hippocampal lysate (H), membrane (SPM) and vesicular (LP2) fractions. N=11. *-\*\*\**P*≤0.0337 (Kruskal-Wallis test; F=27.08, *P*<0.0001; Dunn’s post-hoc test). (**M**). Representative widefield images of KI and WT hippocampal neurons in primary dissociated culture labeled with α-GluA2 live (left), after fixation (center) or after fixation and permeabilization (right). Scale bars, 20 µm. (**N**). Normalized fluorescence intensity of surface or whole-cell GluA2. N≥18. *P*≥0.2649 (Mann-Whitney U-tests). (**O**). Representative tiled widefield images of 50 µm frontal sections from KI or WT brains labeled with α-GluA2. Scale bars, 500 µm. (**P**). Fluorescence intensity of GluA2 in KI or WT sections, normalized to secondary-only control. N=4. *P*≥0.1682 (unpaired T-tests). Error bars, SEM. See also Supplementary Figures S11-S15.

We then used dissociated primary hippocampal cultures and frontal brain sections from KI and WT mice to perform immunological characterization of GluA2 expression and localization. We found no difference in the amount of GluA2 at the neuronal surface in live-labeled or fixed-unpermeabilized neurons in culture, or in the total GluA2 content of fixed-permeabilized neurons (Figure 4M-N) or whole brain sections (Figure 4O-P, Figure S15). Taken together, these data confirm normal GluA2 expression and subcellular localization in the AP-GluA2 KI model.

### Crosslinking of biotinylated AP-GluA2 by NeutrAvidin precludes LTP in the hippocampal CA1 region

We next evaluated the impact of AP-GluA2 KI on AMPAR channel function and synaptic organization in adult brain circuits using whole-cell voltage clamp and field electrophysiological recordings in acute hippocampal slices. AMPAR rectification, NMDA/AMPA ratio, excitatory/inhibitory balance, spontaneous excitatory and inhibitory events, and input/output curves were all indistinguishable between KI and WT preparations (Figure 5A-O). This is in line with our previous characterization of the biophysical properties of AP-tagged GluA2 subunits (Penn et al., 2017), and indicates that AP-GluA2-containing AMPAR are fully functional and that the genetic modification of GluA2 does not impact basal synaptic or network physiology.

**Figure 5:**
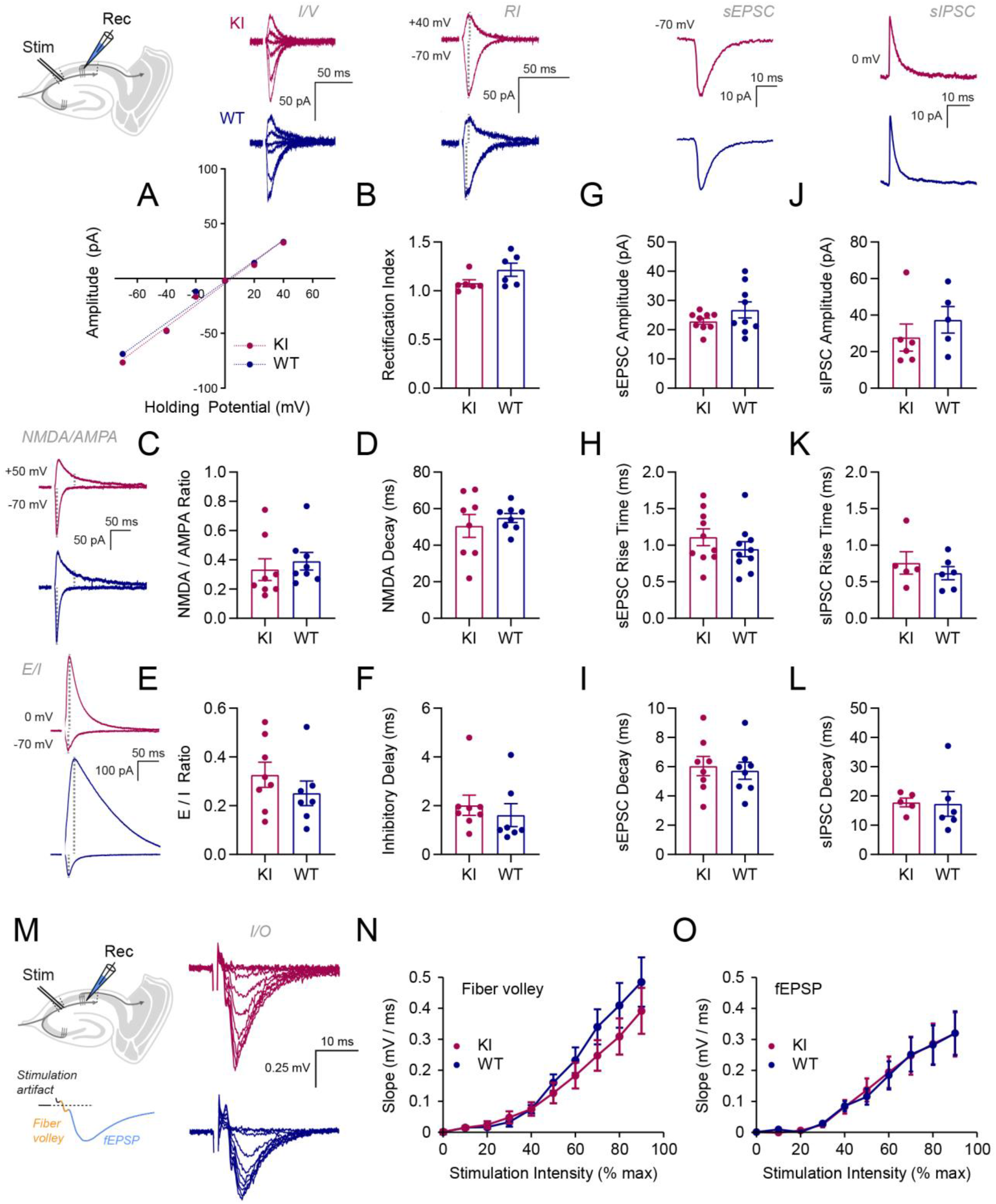
Electrophysiological characterization of AP-GluA2 KI. (**A-L**). Whole-cell voltage-clamp electrophysiological characterization of AMPAR function and synaptic physiology at CA3-CA1 Schaeffer collaterals in acute slices from adult AP-GluA2 KI or WT mice. Typical current responses in CA1 pyramidal neurons and holding potentials are shown in inserts above or left of the relevant panels (KI in red; WT in blue). Representative current-voltage (I/V) plot of synaptic AMPA current amplitude (**A**) and rectification index (RI) measured in CA1 pyramidal neurons (**B**). N=6. *P*=0.1320 (Mann-Whitney U-test). Ratio of synaptic NMDA and AMPA currents (**C**) and NMDA current decay time (**D**). N=8. *P*=0.1605 (Mann-Whitney U-test), *P*=0.5351 (unpaired T-test). Feed-forward inhibition evoked by stimulation of Schaeffer collaterals was used to assess the excitation-inhibition balance (E/I) of hippocampal circuits (**E**) and delay of inhibitory current onset (**F**). N≥7. *P*=0.3143 (unpaired T-test), *P*=0.2671 (Mann-Whitney U-test). Characterization of the amplitude (**G**,**J**), rise time (**H**,**K**) and decay time (**I**,**L**) of spontaneous excitatory (**G-I**) and inhibitory (**J-L**) postsynaptic currents (sEPSC/sIPSC). N≥5. *P≥*0.2134 (unpaired T-test or Mann-Whitney U-test). (**M-O**). Functional impact of AP-GluA2 KI was assessed using extracellular field recordings of presynaptic fiber volleys (FV) and excitatory postsynaptic potentials (fEPSP) resulting from Schaeffer collateral stimulations of increasing intensity. Typical responses for stimulations between 10-90% of maximal intensity are shown in (**M**). Input/output (I/O) curves of FV (**N**) and fEPSP (**O**) responses to stimulations of increasing intensity recorded in the CA1 region of acute slices from adult KI and WT mice. N≥14. *P*≥0.0539 (unpaired T-test or Mann-Whitney U-test). Error bars, SEM.

To determine if long-term plasticity is intact at CA3 to CA1 Schaeffer collaterals in AP-GluA2 KI mice, we used high frequency stimulation (HFS; 3×1s at 100 Hz) to induce LTP at CA3-CA1 synapses in acute slices from KI and WT mice. In both preparations HFS application was followed by a significant increase in the excitatory postsynaptic potential/fiber volley (fEPSP/FV) slope ratio, with a comparable increase in the synaptic response (Figure 6A,C-D), indicating that appropriate synaptic plasticity is maintained with the AP-GluA2 genetic modification. To achieve *in vivo* biotinylation of AP-GluA2, we performed stereotaxic injection of AAVs encoding BirA^ER^-eGFP or eGFP control into the CA1 region of AP-GluA2 KI mice. Biotin supplementation was achieved by five consecutive days of intraperitoneal (IP) injections before the preparation of acute slices, where NA was applied by preincubation of the slices with 100 nM NA before transfer to the recording chamber, and field recordings were made from areas with a high density of eGFP-expressing neurons in ACSF containing 10 pM NA to immobilize newly exocytosed receptors (Figure 6B) (Penn et al., 2017). In slices from BirA^ER^ AAV-injected AP-GluA2 mice, the level of LTP expression after HFS was almost completely abolished when bAP-GluA2 were crosslinked by NA, while it remained normal in slices from eGFP AAV-injected mice (Figure 6E-F). Of note, we observed an initial post-tetanic potentiation, most likely of pre-synaptic origin, which remained unaffected in the presence of NA and bAP-GluA2. These results demonstrate that NA-mediated immobilization of bAP-GluA2-containing AMPAR is an efficient tool to control the expression of activity-dependent synaptic plasticity without impacting basal synaptic network function.

**Figure 6:**
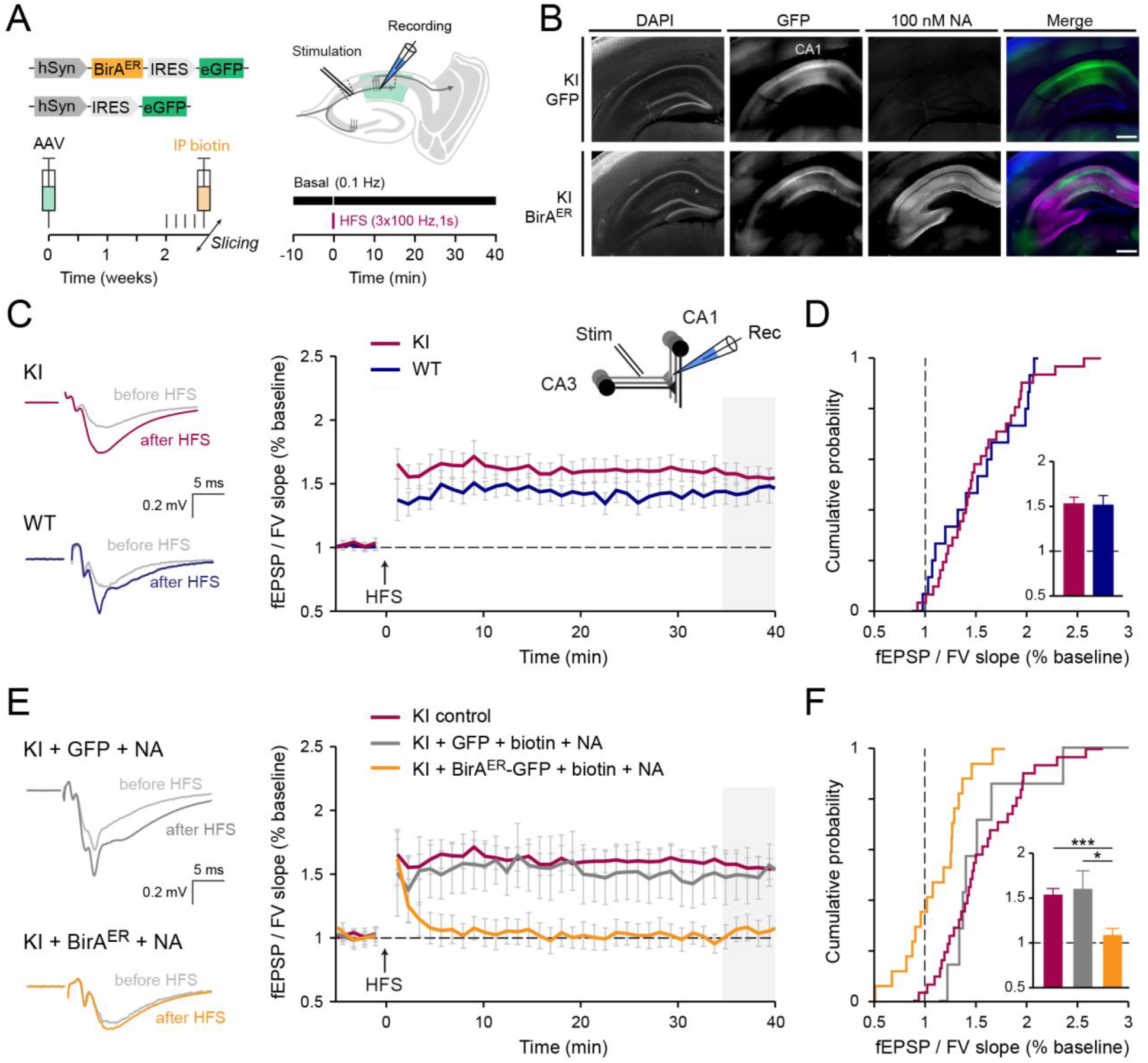
AMPAR immobilization by NA crosslinking precludes LTP. (**A**). Overview of the experimental preparation, AAV stereotaxic injection in the dorsal hippocampus followed by IP biotin injection (left), and field recordings of LTP in CA1 evoked by high frequency stimulations (HFS; 3x 1s trains, 100 Hz) of Schaeffer collaterals in acute slices (right). (**B**). Representative tiled widefield images of 300 µm frontal brain sections from KI mice injected with eGFP control or BirA^ER^-eGFP AAV, live-labeled with 100 nM NA conjugated to DyLight 633. Scale bars, 500 µm. (**C-D**). HFS-induced LTP in acute slices from KI and WT mice (KI in red; WT in blue), representative voltage traces and summary plots (**C**), and cumulative histograms (**D**) of mean normalized fEPSP/FV slope. N≥16. *P*=0.8580 (unpaired T-test). Statistical comparison of LTP was 35-40 min after HFS induction (shaded grey). (**E**-**F**). HFS-induced LTP in acute slices from KI mice injected in CA1 with eGFP control or BirA^ER^-eGFP AAV (eGFP in grey; BirA^ER^ in orange), slices were incubated with 100 nM NA then continuously perfused with 10 pM NA, as in (**C**,**D**). N≥8. *-\*\*\**P*≤0.0280 (Kruskal-Wallis test; F=14.94, *P*=0.0006; Dunn’s post-hoc test). Error bars, SEM.

### Crosslinking of biotinylated AP-GluA2 by NeutrAvidin prevents the formation of contextual fear memory

LTP mechanisms in the dorsal hippocampus have been linked to memory acquisition *in vivo* (Whitlock et al., 2006), and we have previously reported that antibody-mediated crosslinking of AMPAR surface diffusion impaired the formation of contextual fear memories in mice (Penn et al., 2017). Therefore, we reasoned that crosslinking of bAP-GluA2 with NA would afford target-specific control of memory formation and fear behavior in AP-GluA2 KI mice upon the expression of BirA^ER^. To this end, we performed bilateral stereotaxic injection of BirA^ER^-eGFP or eGFP AAVs in the CA1 region of AP-GluA2 KI mice, and implanted guide cannulas to allow infusion of NA into the dorsal hippocampus *in vivo* (Figure 7A-B). Compared to the control conditions of BirA^ER^ + saline infusion or eGFP + NA infusion, mice injected with BirA^ER^ and infused with NA exhibited significantly reduced levels of freezing when re-exposed to the fear-conditioned context (Figure 7C-E; vi in Figure 7A). All the groups exhibited appropriate hippocampus-independent cued fear memory (Figure 7C,F-G; v in Figure 7A). This demonstrates that *in vivo* immobilization of surface diffusing AMPAR with NA offers a new approach to control associative memory and opens to target-specific control of behaviorally-relevant synaptic plasticity expression *in vivo*.

**Figure 7:**
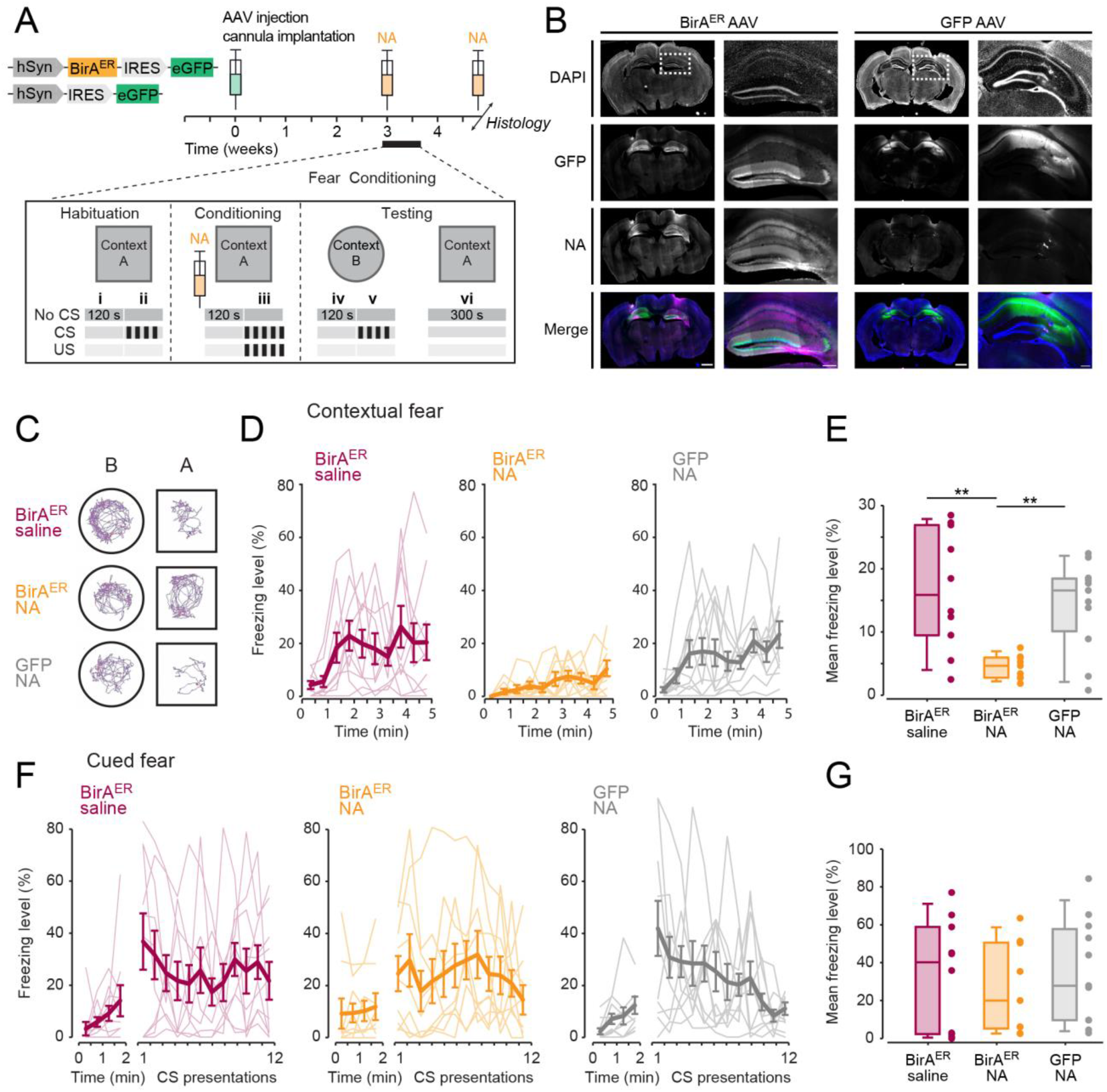
AMPAR immobilization by NA crosslinking blocks formation of contextual fear memories. (**A**). Overview of the experimental design, stereotaxic injection of BirA^ER^-eGFP or eGFP control AAV in the dorsal hippocampus and surgical implantation of the cannula used to deliver NA before fear acquisition and post-mortem histology to verify infusion sites (i-ii: Basal freezing; iii: Fear acquisition; iv: Contextual fear control; v: Cued fear expression; vi: Contextual fear expression). (**B**). Representative tiled widefield images of 60 µm frontal brain sections from KI mice injected with BirA^ER^-eGFP or eGFP control AAV, and infused with 8.33 µM NA conjugated to Texas Red by implanted cannula. Scale bars, 1000 and 250 µm. (**C**). Representative traces of mouse movements during the first two min of exposure to context A and B. (**D-E**). Time course of freezing behavior expression (**D**) and mean freezing levels (**E**) observed during contextual fear memory testing. N≥9. \*\**P*≤0.0074 (Welch’s ANOVA, F=15.64, *P*=0.0003; Dunnett’s post-hoc test). (**F**-**G**). Freezing behavior observed upon CS presentation during cued fear memory testing, as in (**D**-**E**). N≥9. *P*≥0.7577 (one-way ANOVA, F=0.2724, *P*=0.7636; Tukey’s post-hoc test).

## DISCUSSION

Advancements in understanding the organization and function of the brain are constrained by the limitations of currently available techniques, where experimental access to appropriate spatiotemporal resolutions, endogenous proteins, and opportunities to assay synaptic and neuronal function within complex integrated circuits remain formidable challenges. Bridging this gap requires parallel developments in high-resolution imaging methods and molecular tools to visualize and functionalize proteins of interest. In this study we have developed and characterized a new mouse model system and an associated molecular toolkit, where genetic KI of the AP tag and target-specific regulation of BirA expression allows the surface trafficking dynamics of endogenous AMPAR to be monitored and manipulated with avidin probes that efficiently access confined synaptic domains in thick biological tissues. Importantly, by tuning BirA expression, the resolution of this system can be scaled from the study of single molecules in individual neurons to integrated circuits in behaving animals. This opportunity for sparse, target-specific labeling of endogenous surface AMPAR offers a significant advantage for high-resolution synaptic imaging applications in tissue preparations, which would otherwise be obscured by the high signal density in constitutive genetic KI models. We furthermore anticipate that this technology will be readily adaptable to studying and controlling the nanoscale organization and trafficking dynamics of most cell surface proteins, which can now be accomplished by AP tag functionalization of endogenous proteins with relative ease thanks to CRISPR-Cas9 mediated genome editing of animal models or experimental preparations (Doudna and Charpentier, 2014; Fang et al., 2021; Heidenreich and Zhang, 2016; Nishiyama et al., 2017). It is particularly interesting to note the emerging roles for the lateral diffusion of synaptic adhesion molecules, presynaptic voltage gated calcium channels, and astrocytic glutamate transporters in shaping synapse assembly, function, and plasticity (Chamma et al., 2020; Murphy-Royal et al., 2015; Schneider et al., 2015). Together, these features promise to open new avenues of investigation compatible with a wide range of experimental techniques and biological research questions.

Here, we demonstrate that genetically modified AP-GluA2 subunits exhibit normal protein expression, localization, and function, and that AP-GluA2 KI mice exhibit normal synaptic physiology, circuit function, and behavior. Notably, a recently generated SEP-GluA1 KI mouse line was found to exhibit significant reductions in GluA1 protein expression and synaptic localization, with compensation by upscaling of GluA2/3 (Graves et al., 2020). This suggests that genetic KI of small sequences such as the AP tag is an advantageous strategy for functionalizing AMPAR subunits that better conserves normal expression patterns and synaptic composition. One puzzling observation in the AP-GluA2 KI that is at present difficult to explain is the slower migration and unusual banding pattern of AP-GluA2 in Western blots. We could not find evidence for a role of post-translational modification in this migration and banding pattern. However, after extensive characterization by subcellular fractionation and immunoreactivity assays, we propose as the most parsimonious explanation that GluA2 migration is altered by the presence of charged residues in the AP-TEV KI sequence, and that the AP tag is proteolytically degraded within intracellular compartments, but remains intact in the biogenesis pathway and on the cell surface. As naturally biotinylated proteins are largely absent from cell surface membranes (Chamma et al., 2016; de Boer et al., 2003), we found avidin labeling to be highly specific to bAP-GluA2 and tightly controlled by the expression or application of BirA. One limitation of this approach is the opportunity to quantify absolute receptor numbers at the synapse, as a proportion of AP-GluA2 at the neuronal membrane may remain unbiotinylated and therefore undetectable by avidin probes. It is worth noting, however, that this type of constraint exists for all protein labeling or genetic tagging strategies (Choquet et al., 2021). Together with the remarkably high catalytic efficiency of BirA (Beckett et al., 1999), our observations that AP-GluA2 biotinylation saturated over time with chronic and acute BirA approaches (Figure 1E,I) and that non-biotinylated AP-GluA2 were largely confined to intracellular compartments (Figure 4I-J) suggest that the amount of unbiotinylated AP-GluA2 at the neuronal surface is minor.

The LLSM imaging modality permits high-resolution, high-speed imaging with low photobleaching and photoxicity, which has enabled nanoscale imaging of fast dynamic processes in live tissues and organisms (Chen et al., 2014), and permitted new insights into the morphological organization of the brain by coupling expansion microscopy with the rapid large-volume imaging capacity of LLSM (Gao et al., 2019). Our development of a photostimulation module significantly advances the functionality of LLSM setups by offering new opportunities to perform all-optical physiological studies in live tissue preparations with enhanced 4D spatiotemporal resolution (Ducros et al., 2019). The combination of LLSM-FRAP and mSA or NA labeling of bAP-GluA2 enabled characterization of the mobility/immobilization dynamics of endogenous AMPAR in an integrated brain slice preparation. Whereas previous studies characterizing AMPAR mobility dynamics in hippocampal slices or *in vivo* have mostly used overexpression of SEP-tagged AMPAR subunits (Chen et al., 2021; Heine et al., 2008; Makino and Malinow, 2009), few have accomplished measurement of endogenous AMPAR, with the exception of chemical labeling of native AMPAR complexes in acute slices (Wakayama et al., 2017). Importantly, previous measurements of synaptic AMPAR mobile fractions using SEP overexpression approaches ranged from ∼30-100%, and are generally higher than what we have found in the present study, or what has been found on endogenous AMPAR in cultured neurons (Choquet, 2018; Fang et al., 2021). This lends support to the notion that measurements of AMPAR surface diffusion dynamics are impacted by experimental approaches that manipulate the content of AMPAR surface pools through overexpression, and the importance of measuring the properties of receptors expressed at endogenous levels. Tetravalent crosslinking of bAP-GluA2 by NA application efficiently decreased AMPAR surface mobility at synaptic sites, while extrasynaptic AMPAR remained partly mobile. We suspect that some extrasynaptic receptors escape crosslinking due to the low density on the dendritic shaft, or that small clusters may remain mobile if none of the components are bound to stable anchoring structures, as in the PSD. This likely explains why AMPAR internalization, which occurs at extrasynaptic sites (Blanpied et al., 2002; Petrini et al., 2009), was not affected by AMPAR crosslinking in our measurements (Figure S8-S9).

We urgently need new strategies to progress on understanding the relationship between synaptic plasticity and neuronal network rearrangements induced by memory formation. Several open questions remain, such as the importance of synaptic plasticity mechanisms for encoding, consolidation and retrieval of memories, as well as the role of specific sleep phase oscillations in reactivating and selecting inputs potentiated during memory encoding (Humeau and Choquet, 2019; Tononi and Cirelli, 2014). As the vast majority of excitatory synapses onto pyramidal cells express postsynaptic LTP using GluA2-containing AMPAR (Lu et al., 2009; Nicoll, 2017; Schwenk et al., 2014), we expect that the spatiotemporal control of AMPAR mobility *in vivo* afforded by this experimental model will allow progress in establishing the causality link between synaptic plasticity and memory dynamics. The future development of molecular strategies to control cell-type specific or activity-dependent BirA expression holds great potential to answer as-yet inaccessible questions regarding the role of synaptic plasticity for memory formation in discrete cell populations. For example, enhancer-promoter sequences may be useful to restrict AAV-mediated BirA expression to defined cell types (de Leeuw et al., 2016), and AAV-PHP.eB capsids for non-invasive gene delivery to the CNS may allow for BirA transduction of distinct cell populations that are broadly distributed throughout the brain (Chan et al., 2017). Furthermore, coupling synaptic plasticity blockade with recent techniques to restrict BirA expression to neurons activated during specific behavioral modalities (e.g. FliCRE, Cal-light) would be of great interest to test the importance of a neuron’s activity profile in its future involvement in memory engrams or behaviorally relevant neuronal ensembles (Kim et al., 2020; Lee et al., 2017). While the present crosslinking strategy is highly efficient for blocking synaptic plasticity at most excitatory synapses, two limitations must be noted. Firstly, plasticity must remain unaffected at synapses that express presynaptic LTP, such as hippocampal mossy fibers (Nicoll and Malenka, 1995; Zalutsky and Nicoll, 1990). Secondly, a degree of insensitivity must also exist at excitatory contacts onto interneurons, which have been shown to express synaptic plasticity upon learning and often contain GluA1 homomers (Cummings and Clem, 2020; Dupret et al., 2013; Lucas et al., 2016). Transposing this strategy to the development of AP-GluA1 KI mice should be useful for the study of AMPAR mobility and synaptic plasticity at these excitatory synapses onto interneurons.

In conclusion, this KI mouse model and molecular toolkit represents a new genetic labeling strategy that enables target-specific measurement and control of the mobility dynamics of endogenous cell surface proteins in integrated experimental systems. For studying AMPAR dynamics, this model opens new possibilities to explore as-yet unanswered questions regarding the molecular links between receptor surface mobility, synaptic plasticity, and behavioral adaptation. With the CRISPR-Cas9 revolution in targeted genome engineering, this methodology can be adapted for the study of a large variety of synaptic proteins, with broad implications for advancing our understanding of brain organization and function.

## ACKNOWLEDGEMENTS

This work was supported by European Research Council grants to DC (ERC; grants ADOS 339541 and Dyn-Syn-Mem 787340), a Fondation Recherche Médicale grant to YH (FRM; grant DEQ20180339189 AMPA-MO-CO), an Agence Nationale de la Recherche grant to MS (ANR; grant OptoXL ANR-16-CE16-0026), and the Conseil Régional de Nouvelle Aquitaine. AG was funded by Canadian Institutes of Health Research (CIHR; grant 158090) and University of Bordeaux Initiative of Excellence (IdEx) postdoctoral fellowships.

This work was supported by the Bordeaux Neurocampus core facilities (LabEx BRAIN; grant ANR-10-LABX-43), including the cell biology, EOPS and PIV-EXPE facilities of the IINS, the biochemistry and biophysics platform and animal genotyping facility of Neurocentre Magendie (INSERM), and the AAV production platform of the IMN. The microscopy was done at the Bordeaux Imaging Centre, a service unit of CNRS-INSERM and Bordeaux University, a member of the national infrastructure France BioImaging supported by the French National Research Agency (grant ANR-10-INBS-04). The authors wish to thank E. Verdier, N. Retailleau, N. Chevrier, Y. Rufin, C. Lemoigne, H. El Oussini-Ben Chaabane, T.-A. Vernoy, G. Dabee, E. Normand, P. Costet, M. Dehors, C. Martin, C. Castelli, R. Sterling, E. Herzog, J. Teillon, M. Malivert, and S. Marais for their contributions to this work.

## RESOURCE AND MATERIALS AVAILABILITY STATEMENT

All datasets generated or analyzed during this study are included in the published article and its associated supplementary information file. Requests for further information or resource and reagent availability should be made to Daniel Choquet.

## AUTHOR CONTRIBUTIONS

Experimental conception and design: AG, CB, YH, DC. Manuscript preparation: AG. All authors discussed the results and edited the manuscript. LLSM and FRAP: AG, MD. Molecular biology and biochemistry: AG, CB, SD, MS. Biotinylation and TEV experiments: AG, AN. Immunological experiments: AG. dSTORM and uPAINT: HZ, EH. TEM: MFM. Electrophysiology: UF, FL, YH. Animal surgeries and behavior experiments: ALSA, UF.

## METHODS

### Animals

The AP-GluA2 KI model was generated by PHENOMIN Mouse Clinical Institute (Strasbourg, France) using CRISPR-Cas9 genome editing of *Gria2* on C57BL/6N embryos. This line originated from a male F0 founder, which was backcrossed on a C57BL/6J background at the PIV-EOPS facility of the IINS to generate the AP-GluA2 KI mouse line used in this study (B6J-Gria2^em1(AP-TEV)Ics/Iins^ N2). Genotyping was performed by PCR assay on tail biopsies by the genotyping facility of Neurocentre Magendie (Bordeaux Neurocampus). See also Supplementary Figure S1. Primers used for genotyping KI and WT animals are shown in Table S1. We used littermate or age-matched KI and WT control animals from the B6J-Gria2^em1(AP-TEV)Ics/Iins^ N2 line, as appropriate. The KI mutation did not impact animal weight, size, growth or fertility. We used the SHIRPA protocol (Hatcher et al., 2001; Rogers et al., 1997) to assess the behavioral phenotype of AP-GluA2 KI mice, and found no significant differences compared to WT littermates of 6-24 week old mice (Figure S16). Animals were housed under a 12 hour light/dark cycle with unrestricted access to food and water. Dissociated hippocampal neuron cultures were prepared from P0 male or female mice and organotypic hippocampal slice cultures were prepared from P5-8 male or female mice. Histology, biochemistry, surgical manipulations, acute slice recordings, and behavior experiments were performed on 4-12 week old male mice. All experiments were performed in accordance with the European guidelines for the care and use of laboratory animals, and the guidelines issued by the University of Bordeaux animal experimental committee (CE50; Animal facilities authorizations A5012009, A3306940 and A33063941; Ethical project authorizations 20778-2019021913051936 and 18507-201901118522837). Every effort was made to minimize the numbers and suffering of experimental animals.

### Molecular Biology

The BirA^ER^ coding sequence was a gift from Alice Ting (Howarth et al., 2005). BirA^ER^ was cloned upstream (5’) of an encephalomyocarditis virus (EMCV) internal ribosome entry site (IRES) sequence, and the eGFP reporter or Cre recombinase was cloned downstream (3’) of the IRES sequence, such that the start codon of the BirA^ER^ signal sequence corresponded to the 11^th^ ATG of the IRES sequence. An IgK leader sequence and hemagglutinin (HA) epitope tag was added to the 5’ end of BirA^ER^. BirA^ER^-eGPP, BirA^ER^-Cre and eGFP constructs were cloned into the multiple cloning site of the pAAV vector (AAV_pSyn backbone). A CAG promoter was used in plasmids prepared for SCE, and a hSyn promoter was used in plasmids prepared for the synthesis of AAV particles. Plasmids were prepared using the ZymoPURE plasmid MaxiPrep kit (Zymo Research, ZD4203). All constructs were verified by restriction enzyme digestion and Sanger DNA sequencing. AAV serotype 1 and 9 preps were produced by the viral core facilities of the Bordeaux Neurocampus IMN, Charité Universitätsmedizin Berlin, or ordered from Addgene. pENN.AAV.hSyn.Cre.WPRE.hGH was a gift from James M. Wilson (Addgene plasmid # 105553; Addgene viral prep # 105553-AAV9; http://n2t.net/addgene:105553; RRID:Addgene_105553). pAAV.synP.DIO.EGFP.WPRE.hGH was a gift from Ian Wickersham (Addgene plasmid # 100043; Addgene viral prep # 100043-AAV9; http://n2t.net/addgene:100043; RRID:Addgene_100043). Viral titers were between 3.05×10^13^ – 5.77×10^14^ GCP/mL.

### Primary Dissociated Neuron Cultures

Banker cultures of hippocampal neurons from P0 mice were prepared as previously described (Kaech and Banker, 2006), with modifications. Briefly, pups were anesthetized on ice and sacrificed by decapitation. Dissected hippocampi were treated with papain for 20 min at 37°C, then dissociated in Hibernate-A medium (Gibco, A1247501). Dissociated neurons were plated at a density of 500,000-600,000 cells per 60-mm dish on poly-D-lysine and laminin (Sigma-Aldrich, P6407, 11243217001) pre-coated 18 mm 1.5H coverslips (Marienfeld-Superior, 0117580) in Neurobasal A medium supplemented with 10% heat-inactivated horse serum, 0.5 mM GlutaMAX and B-27 Plus (Gibco, 26050088, 12349015, 35050061, A3582801). After 30 min, coverslips were rinsed with Neurobasal A medium supplemented with 0.5 mM GlutaMAX and B-27 Plus, then flipped onto an astrocyte feeder layer. Neurons were maintained at 37°C with 5% CO_2_ in. 2 µM Ara-C was added after 72 h to stop glial proliferation. Astrocyte feeder layers were prepared two weeks in advance from P0 WT mice, plated at a density of 50,000 cells on poly-L-lysine (Sigma-Aldrich, P2636) pre-coated 60-mm dishes, and maintained at 37°C with 5% CO_2_ in MEM supplemented with 4.5 g/L glucose, 2 mM GlutaMAX and 10% heat-inactivated horse serum (Gibco, 26050088). Cultured neurons were transduced with BirA^ER^-eGFP or eGFP AAV1 at DIV 3-7 by incubating coverslips overnight in 12-well plates with 0.5 mL pre-conditioned Neurobasal A media containing viruses at a MOI of 10,000-20,000. Coverslips were then returned to the 60-mm dishes and maintained with 10 µM D-biotin supplementation (Sigma-Aldrich, B4639) for 2-3 weeks. 20-30% of the media was changed 1-2 times per week.

### Organotypic Slice Cultures

Organotypic hippocampal slice cultures from P5-8 mice were prepared as previously described (Stoppini et al., 1991), with modifications. Briefly, pups were anesthetized on ice and sacrificed by decapitation. Hippocampi were dissected in media containing (in mM) 10 D-Glucose, 4 KCl, 26 NaHCO_3_, 234 Sucrose, 5 MgCl_2_, 1 CaCl_2_, and 1 Phenol Red, equilibrated for ∼5 min with carbogen (5% CO_2_ and 95% O_2_). 300 µm transverse slices were cut with a tissue chopper (WPI McIlwain), then positioned on 0.45 µm Durapore membranes on Millicell culture inserts (Millipore, FHLC01300, PICM0RG50) in 6-well plates. Slices were maintained at 35°C with 5% CO_2_ in MEM containing 20% heat-inactivated horse serum, 1 mg/L insulin, and (in mM) 30 HEPES, 13 D-glucose, 5.2 NaHCO_3_, 1 L-glutamine, 0.25 ascorbate, 1 CaCl_2_, 2 MgSO_4_ (Sigma-Aldrich, M4642, I0516). The medium was replaced every 2-3 days.

Slices were transduced with BirA^ER^-eGFP, eGFP, BirA^ER^-Cre + FLEx eGFP or Cre + FLEx eGFP AAV9 at DIV1 by microinjection of the virus (3-5 pulses, 30 ms, 10 PSI) into the CA1 pyramidal cell layer with glass microelectrodes (∼1-2 MΩ; Science Products, GB150F-10P) (Wiegert et al., 2017b). BirA^ER^-eGFP, eGFP, and FLEx eGFP viruses were used at a dilution of 1:10 – 1:20 in 1X PBS. BirA^ER^-Cre or Cre were used with FLEx eGFP at a dilution of 1:5,000 – 1:20,000. Samples were maintained in media on culture inserts and visualized under a stereomicroscope (Nikon SMZ 745T, Lumenera Infinity1). The microelectrode was positioned with a micromanipulator (Scientifica PatchStar) and viruses were injected using a Picospritzer (Parker Picospritzer III).

CA1 pyramidal neurons were electroporated at DIV3 with glass microelectrodes (∼4-6 MΩ) filled with an internal solution containing (in mM) 135 K-gluconate, 0.2 EGTA, 10 HEPES, 4 MgCl_2_, 4 Na_2_-ATP, 0.4 Na-GTP, 10 Na_2_-phosphocreatine, 3 ascorbate (pH 7.2, 290 mOsm) and 13 ng/µL BirAER-eGFP or eGFP plasmids (Wiegert et al., 2017a). Samples were maintained in pre-warmed Tyrode’s solution containing (in mM) 10 D-glucose, 10 HEPES, 120 NaCl, 3.5 KCl, 2 MgCl_2_, 2 CaCl_2_, 2 NaHCO_3_, 1 Na-Pyruvate (pH 7.3, 300 mOsm), on an upright microscope (Nikon Eclipse FN1, DS-Fi3), the microelectrode was positioned with a micromanipulator (Scientifica PatchStar), and cells were electroporated after the formation of a loose seal (4 x 25 ms pulses, 1 Hz, −2.5V; NPI ISO-STIM 01D, Multi Channel Systems STG 4002, Voltcraft FPS-1132). After AAV transduction or SCE, slices were maintained in culture media supplemented with 10 µM biotin.

### Biotinylated AP-GluA2 Labeling and TEV experiments

mSA was produced and conjugated to STAR 635P (Abberior, ST635P) using NHS-ester-activated fluorophore coupling as previously described (Chamma et al., 2017). NA was conjugated to STAR 635P as above (ThermoFisher, 31000), or NA-dye conjugates were purchased from commercial suppliers (Abberior STAR 635P, ST635P-0121; Invitrogen DyLight 550 or 633, 84606, 22844). mSA and NA labeling experiments were performed on primary hippocampal cultures at DIV 20-23, or organotypic slices from DIV 12-15, with the exception of the biotinylation time-course experiments, which were performed from DIV 5-23.

For dissociated cultures, coverslips were washed twice for 5 min in Tyrode’s solution containing (in mM) 10 D-glucose, 10 HEPES, 110 NaCl, 5 KCl, 5 MgCl_2_, 2CaCl_2_ (pH 7.4, ∼260 mOsm) with 2% biotin-free BSA (Roth, 0163), then incubated for 5 min with 100 nM NA and washed twice for 5 min with Tyrode-BSA. For TEV proteolytic cleavage, coverslips were washed in Tyrode’s without BSA, incubated for 10 min with 100 U AcTEV (Invitrogen, 12575-015) or vehicle control, then washed twice for 5 min in Tyrode’s. Above steps were performed at 37°C with 5% CO_2_. Cells were fixed for 10 min with 4% PFA-sucrose, washed three times with 1X PBS, incubated for 10 min with 50 mM NH_4_Cl, washed three times with 1X PBS, then rinsed with H_2_O and mounted with Fluoromount-G + DAPI (ThermoFisher, 00-4959-52).

For organotypic slices, samples were washed twice for 10 min in Tyrode’s solution containing (in mM) 10 D-glucose, 10 HEPES, 120 NaCl, 3.5 KCl, 2 MgCl_2_, 2 CaCl_2_, 2 NaHCO_3_, 1 Na-Pyruvate (pH 7.3, 300 mOsm) with 1% biotin-free BSA, then incubated for 20 min in ∼ 30 µL Tyrode-BSA with 100 nM or 400 nM mSA or NA and washed twice for 10 min with Tyrode-BSA. For TEV proteolytic cleavage, slices were washed with Tyrode’s without BSA, incubated for 10 min with 100 U AcTEV (Invitrogen, 12575-015) or vehicle control, then washed twice for 5 min in Tyrode’s. For the AMPAR internalization assay, slices were incubated after mSA or NA labeling in Tyrode’s without BSA, with 1 µM tetrodotoxin citrate (TTX; Tocris, 1069), or treated for 3 min with 30 µM N-Methyl-D-aspartic acid (NMDA) (Sigma-Aldrich, M3262). After 30 min slices were incubated for 10 min with AcTEV or vehicle control, as above. For biotinylation of AP-GluA2 by soluble recombinant BirA, slices were briefly washed with Tyrode’s, then incubated for 5, 15, 30, 60, or 90 min in ∼ 30 µL Tyrode’s with 10 µM biotin-AMP (Jena Bioscience; NU-894-BIO-S) and with or without 0.3 µM recombinant BirA (Sigma-Aldrich; SRP0417) (Howarth and Ting, 2008). Slices were then incubated with NA as above. For two-colour labelling, slices electroporated with BirA^ER^-eGFP at DIV 3 were first incubated with NA-DyLight 550, then incubated for 30 min with 10 µM biotin-AMP and 0.3 µM recombinant BirA, followed by incubation with NA-STAR 635P. Above steps were performed at 35°C with 5% CO_2_. Slices were fixed and mounted as above, except fixation for slices was 2 h at 4°C.

### Biochemistry and Western Blots

Whole brain (minus cerebellum) and isolated hippocampal protein samples were prepared from 6 week old mice. Tissue was homogenized in 5 mL isomolar buffer containing (in mM) 4 HEPES, 320 Sucrose (pH 7.4), with Calbiochem protease and Halt phosphatase inhibitor cocktails (Millipore, 539134; ThermoFisher, 78420). Subcellular fractionation was performed as described previously (De-Smedt-Peyrusse et al., 2018; Whittaker et al., 1964). All steps were performed at 4°C. For the TEV proteolytic cleavage assay, protein samples were incubated with AcTEV or vehicle control for 1 h at 30°C. For the *in vitro* biotinylation assay, protein samples were incubated with 10 µM biotin-AMP with or without 0.3 µM recombinant BirA in 1X PBS with 5 mM MgCl_2_ (pH 8.0) for 1 h at 30°C (Howarth and Ting, 2008). For the deglycosylation assay, protein samples were incubated in glycoprotein denaturing buffer for 10 min at 100°C, then incubated for 2h at 37°C with 1% NP-40, with or without 500 U endoH, 500 U PNGase F, or 40,000 U O-Glycosidase (NEB, P0702S, P0704S, P0733S). Protein concentration was determined with a Pierce BCA assay (ThermoFisher, 23225) before loading 4-15% SDS-PAGE with the PageRuler plus prestained protein ladder (ThermoFisher, 26619). Semi-dry transfers were done for 10 min with Trans-Blot Turbo HMW (Bio-Rad). Membranes were blocked for 1 h with Intercept Blocking Buffer (Li-Cor, 927-70001), and immunoblotted overnight at 4°C with shaking in Intercept Blocking Buffer with the following antibodies: α GluA2 (1 µg/mL; Millipore, MAB397 or Alomone, AGC-005); α GluA1 (1 µg/mL; Neuromab, 75-327); α PSD-95 (1 µg/mL; Neuromab, 75-028); α γ8 (0.1 µg/mL; Frontier Institute, TARPgamma8-RbAf1000); α stargazin (1 µg/mL; CST, 8511), α CaMKII (1 µg/mL; Millipore, 05-532) α β-actin (0.8 µg/mL; Sigma-Aldrich, 5316); α synaptophysin (0.12 µg/mL; Abcam, 32127); α AP tag (1 µg/mL; Kerafast, EGO016). After three washes in a buffer containing (in mM) 25 Tris, 137 NaCl, 2.7 KCl and 0.05% Tween 20, secondary antibodies conjugated to IRDye800CW or IRDye680LT (0.2 µg/mL; LiCor. 926-68020, 926-68021, 926-32211) or StreptAvidin-IRDye800CW (0.2 µg/mL; LiCor, 926-32230) were used for revelation for 1 h at room temperature with shaking in Intercept Blocking Buffer. Blots were imaged with Odyssey FC or CLx scanners and band intensities were analyzed with Image Studio Lite version 5.2 (LiCor). Linescan analysis was performed with Fiji ImageJ 1.53c (NIH).

### Immunocytochemistry and Immunohistochemistry

For live labeling of dissociated cultures at DIV 20-23, coverslips were washed twice for 5 min in Tyrode’s solution with 1% biotin-free BSA, then incubated for 10 min with 15F1 α GluA2 (2 µg/mL), which was provided by Eric Gouaux (Giannone et al., 2010). Coverslips were then washed twice for 5 min with Tyrode-BSA and rinsed once with Tyrode’s without BSA. Above steps were performed at 37°C with 5% CO2. Cells were fixed for 10 min with 4% PFA-sucrose, washed three times with 1X PBS, incubated for 10 min with 50 mM NH4Cl, then washed three times with 1X PBS. Blocking was done for 10 min in 1X PBS with 1% BSA, then cells were incubated for 30 min with the secondary antibody (2 µg/mL; Goat α mouse-Alexa 568, ThermoFisher, A-21124) and washed three times with 1X PBS. For labeling of fixed cells, coverslips were fixed as above, then incubated for 10 min in 1X PBS with 1% BSA, with or without 0.1% Triton X-100 for permeabilization. Blocking was done as above, then coverslips were incubated for 30 min with 15F1 α GluA2 (2 µg/mL), washed three times with 1X PBS, then incubated for 30 min with Goat α mouse-Alexa 568 and washed three times with 1X PBS. Coverslips were rinsed with H_2_O and mounted with Fluoromount-G + DAPI.

For immunohistochemistry, 9 week old mice were anesthetized with Ketamine/Xylazine (130 and 13 mg/kg) before transcardial perfusion with 1X PBS then 4% PFA. Brains were post-fixed overnight in 4% PFA at 4°C, then washed three times with 1X PBS and incubated overnight with 30% sucrose in 1X PBS at 4°C. 50 µm frontal sections were cut in 1X PBS on a vibratome (Leica, VT1200). Floating sections were rinsed with 1xTBS, then permeabilized for 1 h with 0.3% Triton X-100 and 5% NGS (Gibco, PCN5000). Slices were then incubated overnight with or without 15F1 α GluA2 (1 µg/mL) in 1xTBS with 0.1% Triton X-100 and 5% NGS at 4°C. Slices were then washed three times for 10 min with 1xTBS, and incubated for 2 h with Goat α mouse-Alexa 568 (1 µg/mL), then washed three times with 1xTBS. Except for primary antibody incubation above steps were performed at room temperature with shaking. Slices were rinsed with H_2_O, and mounted with Fluoromount-G + DAPI.

### Widefield and Confocal Imaging

Widefield imaging of cultured neurons and organotypic slice samples was done with a DM5000 (Leica) under HC PL Fluotar 5x NA 0.15, HC PL Fluotar 10x NA 0.3, HCX PL Fluotar 20x NA 0.5, or HCX PL APO 63x oil NA 1.40 objectives (Leica). Fluorescence excitation was done with a LED SOLA light (Lumencor), and emission was captured by an ORCA-Flash4.0 V2 camera (Hamamatsu) controlled by MetaMorph software (Molecular Devices). Mosaics were done with a motorized XY stage (Märzhäuser). Brain slices were imaged with a Nanozoomer 2.0HT with fluorescence imaging module (Hamamatsu) using a UPS APO 20x NA 0.75 objective combined to 1.75x lens (Nikon), for a final magnification of 35x. Fluorescence excitation was done with a LX2000 200W mercury lamp, and emission was captured by a TDI-3CCD camera (Hamamatsu). Confocal imaging was done with a TCS SP8 or SP5 (Leica). The SP8 was mounted on a DM6 FS upright stand with HC Plan Fluotar 10x dry NA 0.30, HCX Plan Apo CS 20x multi-immersion NA 0.70, and HC Plan Apo CS2 63x oil NA 1.40 objectives (Leica). The SP8 was equipped with a motorized XY stage and a galvanometric Z stage, 405, 488, 552 and 638 nm laser lines, a conventional scanner (10-1800 Hz), two internal PMT and two internal hybrid detectors for fluorescence detection, and one external PMT for transmission. The SP5 was mounted on a DM6000 upright stand with HCX Plan APO CS 10x dry NA 0.40, HCX Plan Apo CS 20x multi-immersion NA 0.70, HCX Plan Apo CS 40x oil NA 1.25, HCX Plan Apo CS 63x oil NA 1.40 (Leica). The SP5 was equipped with a motorized XY stage and a galvanometric Z stage, 405, 488, 561 and 633 nm laser lines, a conventional scanner (10-2800 Hz), three internal PMT and two internal hybrid detectors for fluorescence detection, and one external PMT for transmission. Imaging parameters were kept the same amongst relevant samples. Image analysis was performed with ImageJ 1.53c (NIH).

### Lattice Light Sheet Microscopy and FRAP Imaging

The LLSM was built according to the technical information provided by the group of Eric Betzig at Janelia Research Campus, Howard Hughes Medical Institute (HHMI), USA (Chen et al., 2014). The LLSM was used under license from Janelia Research Campus, HHMI. The lattice light sheet (LLS) was focused by a custom 28.6x 0.66 NA 3.74 mm excitation objective (EO; Special Optics). Fluorescence was collected with a CFI APO LWD 1.1 NA 25x 2.0 mm detection objective (DO; Nikon) and imaged on a sCMOS ORCA-Flash4.0 V2 camera (Hamamatsu). The annular mask minimum and maximum NAs were 0.44 and 0.55, respectively. We characterized the LLSM optical resolution using 170 nm diameter beads and found values near diffraction limits laterally and better than confocal microscopy axially. The LLSM setup was modified by addition of a photostimulation module (PSM). 30% of the laser combiner output was sent to the PSM by a variable beamsplitter. A mechanical shutter (Uniblitz, VS25) controlled the photostimulation duration. Sub-micron positioning and patterning was achieved by a set a galvanometric mirrors (XYT) (Cambridge Technology, 6215H) that were optically conjugated to the DO back focal plane. The photostimulation beam was magnified to fill the DO back aperture and coupled to the LLSM detection arm by a quadband dichroic mirror (Semrock, Di03-R405/488/561/635). Shutter timing and galvo positioning were controlled by a USB A/D card (National Instruments, 6002), programmed by a user interface written in LabVIEW (National Instruments). The photostimulation beam diameter measured on beads was near diffraction limited (0.43 µm), and sub-micron diameter was maintained in the sample (0.69 µm). See also Supplementary Figure S7. To ensure sub-micrometric targeting of user defined ROIs, the FRAP galvanometric mirrors were calibrated before every experimental session. Briefly, in a homogeneously fluorescent sample a set of 9 points corresponding to 9 different galvo voltages pairs (Vx,Vy) were illuminated. Their positions were detected on LLSM camera images as 9 pixels (Nx,Ny) for each voltages pair (Vx,Vy). An affine transformation matrix was computed to determine the correspondence between camera image pixels and FRAP galvo voltages. A set of 10 randomly selected points were then targeted, and the distance between the targeted and detected spots was measured. The average of these 10 distances provided the mean targeting error, which was less than 0.2 µm for all experimental sessions.

At DIV 10-15, organotypic slices that had been transduced with BirA^ER^-eGFP or BirA^ER^-Cre + FLEx eGFP AAV9 at DIV1 were labeled with 100 nM NA or 400 nM mSA coupled to STAR 635P, as above. Membranes were cut and mounted onto poly-L-lysine pre-coated 5 mm coverslips, then mounted onto the LLSM sample holder with Dow Corning high-vacuum silicone grease (Sigma-Aldrich, Z273554). The imaging chamber was maintained at 34°C and perfused with artificial cerebrospinal fluid (ACSF) containing (in mM) 12.1 D-glucose, 126 NaCl, 2.5 KCl, 2 MgCl_2_, 2 CaCl_2_, 25 NaHCO_3_, 1.25 NaH_2_PO_4_ (300 mOsm), which was equilibrated with carbogen and perfused at a rate of 1.7 mL/min. Slices were maintained in the imaging chamber for up to 2 h. Images were acquired at ∼10-20 µm below the surface of the slice. 3D images of spines and dendrites (typical volume 12.8 x 12.8 x 20 µm^3^) were acquired before FRAP imaging to clearly identify the ROIs and labeling specificity, with 488 nm excitation of eGFP as a volume marker and BirA^ER^ reporter, and 642 nm excitation of mSA-or NA-STAR 635P, the extracellular label for biotinylated AP-GluA2. Illumination power varied depending on the wavelength, sample brightness, labeling intensity, and depth in slice, which ranged from ∼20-200 µW spread over the entire width and thickness of the LLS. Photobleaching during acquisitions was typically less than 20%. Fast sample translation with a piezo stage was used to acquire Z stacks. All images were acquired in dithered square lattice mode. FRAP illumination was 50 ms to a single focal point, the power measured at the back aperture of the DO was below 4 mW and the photobleach efficiency was typically ∼40-80%. Fluorescence recovery was followed by single plane acquisitions (100 ms) in three steps at 10 Hz (15 s), 1 Hz (60 s) and 0.2 Hz (180 s). Recovery curves were corrected for continuous photobleaching and background noise, as previously described (Ashby et al., 2006; Axelrod et al., 1976; Czondor et al., 2012). To analyze the mobile fraction, individual FRAP curves were fit to a nonlinear regression model using one-phase association with a constraint of Y_0_=0 (Prism 8.3.1, GraphPad). 68-88% of FRAP curves were fit successfully and included in the analysis.

### Super-resolution Imaging

For dSTORM imaging, primary neuronal cultures at DIV 21 were live labelled with 15F1 α GluA2 (3.33 µg/mL) for 7 min at 37° C, then fixed as above. Cells were incubated for 30 min with the secondary antibody (2 µg/mL; Goat α mouse-Alexa 647, ThermoFisher, A-21235). dSTORM imaging was performed on a LEICA DMi8 mounted on an anti-vibrational table (TMC), using a HCX PL APO 160x 1.43 NA oil immersion TIRF objective (Leica) and fiber-coupled laser launch (405, 488, 532, 561 and 642 nm) (Roper Scientific). Fluorescence was collected with an Evolve EMCCD camera (Photometrics). Coverslips were mounted on an open Ludin chamber (Life Imaging Services) and 600 µL of imaging buffer was added (Nair et al., 2013). To reduce oxygen exchange during acquisition an 18 mm coverslip was placed on top of the chamber. Image acquisition was driven by Metamorph software (Molecular devices) and one image stack contained 40,000 frames. The size of the region acquired was 512×512 pixels (100 nm/pixel). Keeping the 642 laser intensity constant, the power of the 405 nm laser was increased during acquisition to control the level of single molecules per frame. Multicolor fluorescent microspheres (TetraSpeck, Invitrogen) were used as fiducial markers to register long-term acquisitions and correct for lateral drifts. Intensity-based drift-corrected super-resolution images (25 nm/pixel) were reconstructed using PALM-Tracer software operating as a plugin of MetaMorph (Kechkar et al., 2013).

For uPAINT imaging the microscope was caged and heated to maintain samples at 37°C. Coverslips were mounted on an open Ludin Chamber (Life Imaging Services) and maintained in pre-equilibrated Tyrode’s solution containing (in mM) 12.5 D-glucose, 25 HEPES, 108 NaCl, 5 KCl, 2 MgCl_2_, 2 CaCl_2_ (pH7.4, ∼260 mOsm). Imaging was performed on a Ti-Eclipse microscope equipped with an APO 100x 1.49 NA oil immersion TIRF objective (Nikon) and laser diodes (405, 488, 561 and 642 nm) (Roper Scientific). Fluorescence signal was detected with an Evolve EMCCD camera (Photometrics). To compare diffusion dynamics, KI and WT neurons transduced with BirA^ER^-eGFP or eGFP AAV1 were imaged at DIV 16-21. eGFP+ KI or WT neurons were labeled with a low concentration (∼1 nM) of 15F1 α GluA2 coupled to SeTau 647 (SETA BioMedicals), which was added at to the Ludin Chamber to sparsely and stochastically label endogenous surface GluA2. For crosslink experiments, KI neurons were washed three times with pre-equilibrated Tyrode’s, followed by an incubation with Tyrode’s (control) or 100 nM unconjugated NA for 8 min at 37°C, then washed again. After mounting the sample, eGFP+ neurons were labeled with a low concentration of mSA-STAR 635P (7.7 nM) to sparsely and stochastically label surface biotinylated AP-GluA2. To avoid photo-toxicity the 642 nm laser was activated at low intensity. The TIRF angle was adjusted in order to maximize signal to noise ratio. Image acquisition and control of the microscope were driven by Metamorph (Molecular devices), the acquisition time was 30 ms and 6000-8000 frames at were acquired to record GluA2 lateral diffusion. The diffusion coefficient (D) based on the fit of the mean square displacement curve (MSD) was extracted from the experiments and analyzed with WaveTracer software operating as a plugin of MetaMorph (Kechkar et al., 2013).

### Transmission Electron Microscopy

For TEM imaging, KI organotypic slices transduced with BirA^ER^-eGFP AAV9 were live-labeled at DIV 15 in Tyrode’s with StreptAvidin-FluoroNanoGold, conjugated to Alexa 546 or Alexa 594 and 1.4 nm NanoGold particles (SA-FNG, Nanoprobes Inc). Slices were fixed with 4% PFA-sucrose and 0.2% glutaraldehyde in 1X PBS overnight at 4°C. To confirm SA-FNG staining, slices were imaged in 1X PBS with a SP5 confocal under a HCX IRAPO L 25X water NA 0.95 objective, as above (Leica). SA-FNG and eGFP+ regions were dissected under a stereomicroscope equipped with a NightSea fluorescence system (EMS). The dissected tissue was then incubated in a permeabilizing buffer containing 0.1% Triton X-100 and 0.2% gelatin in 1X PBS, then re-incubated with SA-FNG in permeabilizing buffer to facilitate synaptic labeling. Samples were then subjected to silver enhancement for 5 min (HQ Silver, Nanoprobes Inc) to detect silver-enhanced nanogold particles by conventional EM. Samples were post-fixed with 1% glutaraldehyde in 0.15 M Sorensen’s phosphate buffer (SPB; EMS) for 10 min at 4°C, then incubated with 1% sodium thiosulfate in H_2_O, washed with SPB, and fixed with SPB containing 1% OsO_4_ and 1.5% potassium ferrocyanide for 1 h on ice. Sequential dehydration was performed with 70%, 90%, and 100% ethanol, followed by two incubations with 100% acetone on ice, then samples were embedded for 2 h with 1:1 acetone and epon resin (Embed-812, EMS), followed by 100% epon resin at 60°C for 2 days. TEM imaging was performed in high contrast mode with a H7650 transmission electron microscope (Hitachi) equipped with an Orius SC1000 CCD camera 0(Gatan Inc).

### Stereotaxic Surgery and Cannula Implantation

Mice were positioned in a stereotaxic apparatus (David Kopf Instruments) under continuous isoflurane anaesthesia with a vaporiser system, and treated with intraperitoneal (IP) buprenorphine (0.1 mg/kg) and local lidocaine (0.4 mL/kg, 1% solution). Stereotaxic injections of BirA^ER^-eGPP or eGFP AAV9 were performed on AP-GluA2 KI males aged 4-6 weeks for slice electrophysiology, or aged 6-8 weeks when coupled with cannula implantation for behavioral experiments. The CA1 region was targeted for AAV9 injection with the coordinates (in mm) AP −2; ML ±1.6; DV −1.15. Stainless steel guide cannulae (26 gauge, PlasticsOne) were bilaterally implanted above the hippocampus with the coordinates (in mm) AP −1.6-2.0; ML ±1.9-2.3; DV - 0.3. Guide cannulae were anchored to the skull with dental cement (Super-Bond, Sun Medical Co). After surgery mice recovered from anaesthesia on a heating pad (35°C), and dummy cannulae were inserted into the guides to reduce the risk of infection.

### Acute Slice Preparation and Electrophysiological Recordings

Acute slices were prepared 2-3 weeks following AAV9 stereotaxic injection, and AP-GluA2 KI and WT mice (7-12 weeks) were anaesthetized with isofluorane prior to decapitation. For LTP experiments, 100 µL biotin at 6 mg/mL (pH 7.4) was injected IP daily for 5 days before slicing. Brains were removed and hippocampal parasagittal slices (320-350 μm) were prepared with a vibratome (Leica VT1200s) in an ice-cold solution containing (in mM) 2 KCl, 26 NaHCO_3_, 1.15 NaH_2_PO_4_, 10 Glucose, 220 Sucrose, 0.2 CaCl_2_, 6 MgCl_2_ (pH 7.4, 290–310 mOsm), oxygenated with carbogen. Slices were incubated for 30-45 min at 35°C and then maintained at room temperature before being transferred to the recording chamber perfused at a rate of 1.8 mL/min with ACSF containing (in mM) 119 NaCl, 2.5 KCl, 26 NaHCO_3_, 1 NaH_2_PO_4_, 10 Glucose, 2.5 CaCl_2_, 1.3 MgCl_2_ (300 mOsm), oxygenated with carbogen.

Whole cell patch clamp recordings were made from CA1 pyramidal cells with a borosilicate glass pipette (4-6 MΩ) filled with an internal solution containing (in mM) 125 CsMeSO_3_, 2 MgCl_2_, 1 CaCl_2_, 10 EGTA, 10 HEPES, 4 Na_2_ATP, 0.4 NaGTP and 5 QX-314-Cl. 100 mM spermine was added to the internal solution to quantify the rectification index of AMPA-mediated EPSCs. Spontaneous EPSCs and IPSCs were recorded at a holding potential of −70 and 0 mV, respectively. Spontaneous currents at each potential were averaged and the rise time (20-80%, ms) and decay time constant, determined with a monoexponential fit, were computed. Synaptic responses in CA1 pyramidal cells were elicited by a brief electrical stimulation (0.1 ms, 0.5-2 V) of the Schaffer collaterals with a bipolar electrode placed in the *stratum radiatum* of CA1. For the rectification index (RI), ACSF contained picrotoxin (100 μM) and D-AP5 (25 μM) to block inhibitory and NMDA-mediated synaptic responses. 10 min after entering into whole cell, AMPA-mediated synaptic responses were recorded at different holding potentials ranging from −70 to +40 mV. A linear fit was performed to compute the theoretical EPSC amplitude at +40 mV. The RI is the ratio between the theoretical and experimental value. The NMDA/AMPA ratio was determined in the presence of picrotoxin (100 μM) in the ACSF. The amplitude of the NMDA component 50 ms after the stimulation onset recorded at +50 mV was divided by the amplitude of the AMPA-mediated EPSCs recorded at −70 mV. A monoexponential fit was performed on the EPSC recorded at +50 mV to determine the NMDA EPSC decay (ms). For the excitatory/inhibitory ratio (E/I), synaptic inhibitory responses were recorded at 0 mV, which is the reversal potential for AMPA and NMDA synaptic currents. Synaptic current onsets were determined by using the *Onset* function from NeuroMatic v3.0c (Rothman and Silver, 2018) operating as a plugin of Igor Pro 8.04 (Wavemetrics). Voltage signals were recorded using a MultiClamp 700B amplifier with Clampex 10.7 (Molecular Devices), low-pass filtered at 10 kHz, digitized at 10 kHz and stored on a personal computer for analysis.

For extracellular field recordings, electrical stimulation of Schaffer collaterals and recordings of synaptic responses were made in the CA1 *stratum radiatum* with borosilicate glass pipettes (tip diameter 5-10 µm; resistance 0.2-0.3 MΩ) filled with ACSF. For I/O curves, Schaffer collaterals in the CA1 region were gradually stimulated (0.5 ms, 0-100 µA, 0.1 Hz) and recorded in the *stratum radiatum* of CA1. For LTP recordings, a 10 min baseline of synaptic responses elicited by stimulation at 0.1 Hz was recorded in current clamp mode, then HFS (3x 100 Hz, 1 min) was applied to induce LTP. Synaptic responses at 0.1 Hz were recorded for a further 40 min to observe LTP. For crosslinking experiments, slices were pre-incubated with 100nM NA in ACSF for 20 min. Recordings were then performed in ACSF with 10 pM NA to immobilize newly exocytosed bAP-GluA2 (Penn et al., 2017). The FV and fEPSP slopes were measured and compared to the stimulation intensity. For LTP recordings, the FV and fEPSP responses were normalized to baseline by calculating their slopes before and after HFS, and the normalized ratio is presented. Signals were recorded as above, and analysis was performed with Clampfit 10.7 (Molecular Devices) and SigmaPlot V14 (Systat Software Inc).

### Fear Conditioning

Mice were housed individually in a ventilation area before the start of behavioral training. Two weeks after stereotaxic surgery, the mice were handled for an additional week. To reduce stress of the mice during subsequent experiments, they were trained daily for multiple insertions and removals of dummy cannula. On the first experimental day (day 1) animals were allowed to explore the conditioning context (Context A) for habituation. Both CS+ (30 s duration, consisting of 50 ms tones repeated at 0.9 Hz, tone frequency 7.5 kHz, 80 dB sound pressure level) and CS− (30 s duration, consisting of 50 ms white noise tones repeated at 0.9 Hz, 80 dB sound pressure level) were presented 4 times with a variable interstimulus interval (ISI, 10-30 s). On day 2, infusion cannulae (33 gauge) connected to a 1 μL Hamilton syringe via polyethylene tubing were inserted into the guides, extending beyond the end of the guide cannulae by 2 mm to target the CA1 of the hippocampus. Texas Red-conjugated NA (8.33 µM; Invitrogen, A2665) or saline was infused bilaterally at a rate of 50 nL/min for a total volume of 500 nL per hemisphere, under the control of an automatic pump (Legato 100, Kd Scientific Inc). The injector was maintained for an additional 2 min to allow sufficient diffusion. NA or saline was injected just before presentations of CS/US pairs in context A. The conditioning phase was performed as follows: 5 pairings of CS+ with the US onset coinciding with the CS+ offset (2 s foot shock, 0.6 mA). In all cases, CS− presentations were intermingled with CS+ presentations and ISI was variable over the whole training course. Contextual memory was tested 48 h after conditioning by analyzing the freezing levels during 5 min of context A re-exposure. Cued fear memory was tested 48 h after conditioning by analyzing the freezing levels during the first 120 s and 30 s following the 12 CS+ presentations in Context B. Freezing behavior was recorded in each behavioral session using a FireWire CCD camera (Ugo Basile), connected to automated freezing detection software (ANY-maze, Stoelting Co). Measurements of freezing behavior were done blind and in sessions alternating between experimental groups. Mice within each group were also selected randomly for analysis. At the experimental endpoint, mice were infused with 500 nL NA then anesthetized before transcardial perfusion, as above. 60 µm frontal brain sections were cut on a vibratome (Leica, VT1200) for post mortem verification of bilateral hippocampal AAV9 expression, NA binding, and implanted cannula position.

### Statistical Analysis

All reported results are derived from at least 3 independent experiments to ensure replicability. No statistical test was used to pre-determine sample sizes. For imaging data, analysis was performed blinded to the experimental condition or done with a semi-automated macro to minimize user bias. For electrophysiology and animal behavior experiments, the experimenter was blind to the genotype or AAV identity. Statistical analysis and data plotting was performed with Prism 8.3.1 or 9.1.1 (GraphPad), or SigmaPlot (V14). Data sets were analyzed with the Shapiro-Wilk test for normality, and parametric (*P*>0.05) or non-parametric (*P*<0.05) statistical tests (two-tailed) were performed as appropriate. F test or Bartlett’s test were used to assess equality of variance. Fear conditioning behavioral data was analyzed with the ROUT method to identify outliers from nonlinear regression, and one outlier was removed from the BirA^ER^ + NA test group. Notably, the one outlier in this test group exhibited lower eGFP fluorescence intensity, absence of NA binding upon terminal NA infusion, and a more lateral position of the implanted cannula, suggesting that the lack of effect on fear behavior was due to a lack of AMPAR crosslinking in the CA1 region. Test details and statistical outcomes for all experiments are reported in the relevant figure legends.

## Key Resource Table

**KEY RESOURCES TABLE**
| Reagent or Resource | Source | Identifier |
| --- | --- | --- |
| **Antibodies** |  |  |
| GluA2 (15F1), mouse monoclonal IgG1κ | Provided by E. Gouaux (Giannone et al., 2010) | Clone 15F1 |
| GluA2 | Millipore | Cat # MAB397 |
| GluA2 | Alomone | Cat # AGC-005 |
| GluA1 | Neuromab | Cat # 75-327 |
| PSD-95 | Neuromab | Cat # 75-028 |
| Gamma 8 | Frontier Institute | Cat # TARPgamma8-RbAF1000 |
| Stargazin | CST | Cat # 8511 |
| CaMKII | Millipore | Cat # 05-532 |
| β-Actin | Sigma-Aldrich | Cat # 5316 |
| Synaptophysin | Abcam | Cat # 32127 |
| AP tag | Kerafast | Cat # EGO016 |
| Goat anti-mouse Alexa 568 | ThermoFisher | Cat # A-21124 |
| Goat anti-mouse IRDye680LT | LiCor | Cat # 926-68020 |
| Goat anti-rabbit IRDye680LT | LiCor | Cat # 926-68021 |
| Goat anti-rabbit IRDye800CW | LiCor | Cat # 926-32211 |
| **Biochemical Reagents** |  |  |
| AcTEV | Invitrogen | Cat # 12575-015 |
| Biotin-AMP | Jena Bioscience | Cat # NU-894-BIO-S |
| Recombinant BirA | Sigma-Aldrich | Cat # SRP0417 |
| NMDA | Sigma-Aldrich | Cat # M3262 |
| TTX | Tocris | Cat # 1069 |
| Picrotoxin | Sigma-Aldrich | Cat # P1675 |
| Spermine | Tocris | Cat # 0958 |
| D-AP5 | Tocris | Cat # 0106 |
| endoH | NEB | P0702S |
| PNGase F | NEB | P0704S |
| O-Glycosidase | NEB | P0733S |
| Calbiochem protease inhibitor | Millipore | Cat # 539134 |
| Halt phosphatase inhibitor | ThermoFisher | Cat # 78420 |
| Pierce BCA assay | ThermoFisher | Cat # 23225 |
| PageRuler plus prestained protein ladder | ThermoFisher | Cat # 26619 |
| **Experimental Models** |  |  |
| Mouse: AP-GluA2 KI or WT | Custom | N/A |
| B6J-Gria2<sup>em1(AP-TEV)Ics/lins</sup><br>N2 |  |  |
| **Molecular Probes** |  |  |
| Monomeric StreptAvidin (mSA) | Custom (Chamma et al., 2017) | N/A |
| Unconjugated NeutrAvidin (NA) | ThermoFisher | Cat # 31000 |
| STAR 635P | Abberior | Cat # ST635P |
| NA-STAR 635P | Abberior | Cat # ST635P-0121 |
| NA-DyLight 550 | Invitrogen | Cat # 84606 |
| NA-DyLight 633 | Invitrogen | Cat # 22844 |
| NA-Texas Red | Invitrogen | Cat # A2665 |
| StreptAvidin FluoroNanoGold AlexaFluor 546 (SA-FNG) | Nanoprobes Inc. | Cat # 7416 |
| StreptAvidin FluoroNanoGold AlexaFluor 594 (SA-FNG) | Nanoprobes Inc. | Cat # 7316 |
| StreptAvidin IRDye800CW | LiCor | Cat # 926-32230 |
| **Recombinant DNA and Viruses** |  |  |
| pDisplay: IgK leader HA BirA<sup>ER</sup> | Gift from A. Ting (Howarth and Ting, 2008) | Addgene; Cat # 20856 |
| pAAV: CAG IgK leader HA BirA<sup>ER</sup> IRES eGFP | Custom | N/A |
| pAAV: hSyn IgK leader HA BirA<sup>ER</sup> IRES eGFP | Custom | N/A |
| pAAV: hSyn IgK leader HA BirA<sup>ER</sup> IRES Cre | Custom | N/A |
| pAAV: CAG IRES eGFP | Custom | N/A |
| pAAV: hSyn IRES eGFP | Custom | N/A |
| AAV1/9: hSyn IgK leader HA BirA<sup>ER</sup> IRES eGFP | Custom | N/A |
| AAV1/9: hSyn IRES eGFP | Custom | N/A |
| AAV9: hSyn IgK leader HA BirA<sup>ER</sup> IRES Cre | Custom | N/A |
| AAV9: DIO eGFP | Gift from I. Wickersham | Addgene; Cat # 100043 |
| AAV9: hSyn Cre WPRE hGH | Gift from J. Wilson | Addgene; Cat # 105553 |
| **Software** |  |  |
| Clampex 10.7<br>Clampfit 10.7 | Molecular Devices, LLC | https://www.moleculardevices.com/products/axon-patch-clamp-system/acquisition-and-analysis-software/pclamp-software-suite |
| Igor Pro 8.04 64-bit | Wavemetrics | https://www.wavemetrics.com/products/igorpro |
| NeuroMatic v3.0c | Written by J. Rothman (Rothman and Silver, 2018) | http://www.neuromatic.thinkrandom.com/ |
| ANY-maze 6.3 video tracking | Stoelting Co | https://www.any-maze.com/ |
| Fiji | NIH | https://imagej.net/Fiji |
| MetaMorph | Molecular Devices | https://www.moleculardevices.com/products/cellular-imaging-systems/acquisition-and-analysis-software/metamorph-microscopy#gref |
| WaveTracer, PALM-Tracer | Custom (Kechkar et al., 2013) | https://imagxcell.com/wave-tracer/ |
| LabVIEW | National Instruments | https://www.ni.com/fr-fr/shop/labview.html |
| Lattice Scope | Written by D. Milkie, Betzig team, Janelia Research Campus, HHMI | Custom LLSM acquisition software written in LabVIEW |
| LLSpy V0.4.8 | Written by T. Lambert, Harvard Medical School | https://github.com/tlambert03/LLSpy |
| Prism 8.3.1 and 9.1.1 | GraphPad | https://www.graphpad.com/scientific-software/prism/ |
| SigmaPlot V14 | Systat Software Inc | https://systatsoftware.com/products/sigmaplot/ |
| Adobe Illustrator 22.0.1 | Adobe | www.adobe.com |

## SUPPLEMENTARY EXPERIMENTAL PROCEDURES

**Table S1:**
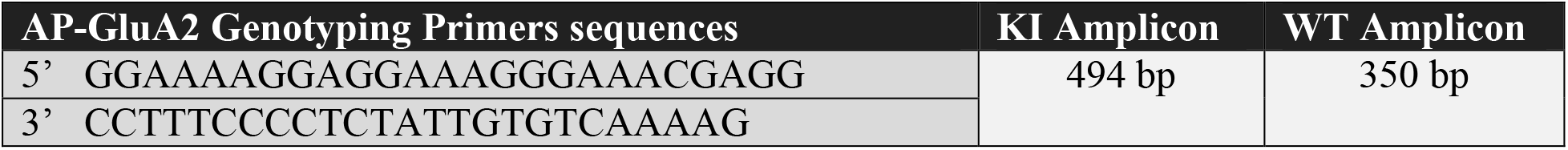
AP-GluA2 KI PCR Genotyping Strategy

### Elevated Plus Maze

As the SHIRPA (SmithKline Beecham, Harwell, Imperial College, Royal London Hospital, phenotype assessment) results suggested a possible effect on anxiety levels of AP-GluA2 KI mice at 14 weeks of age (Supplementary Figure S16A), we complemented this scoring by conducting the elevated plus maze test in adult WT and KI mice (Supplementary Figure S16D). The elevated plus maze apparatus consisted of two opposed open and enclosed arms (50×10×40 cm LWH) and an open central square (10×10 cm LW), elevated 50 cm above the floor. At the beginning of each trial, the mouse was placed at the end of the closed arm. Then animal displacement was assessed for 10 consecutive min using EthoVision XT (Noldus). The percentage of open and closed arm entries were calculated in WT and KI mice at 3 ages (12, 18, 26 weeks).

## SUPPLEMENTARY FIGURES AND FIGURE LEGENDS

**Supplementary Figure S1:**
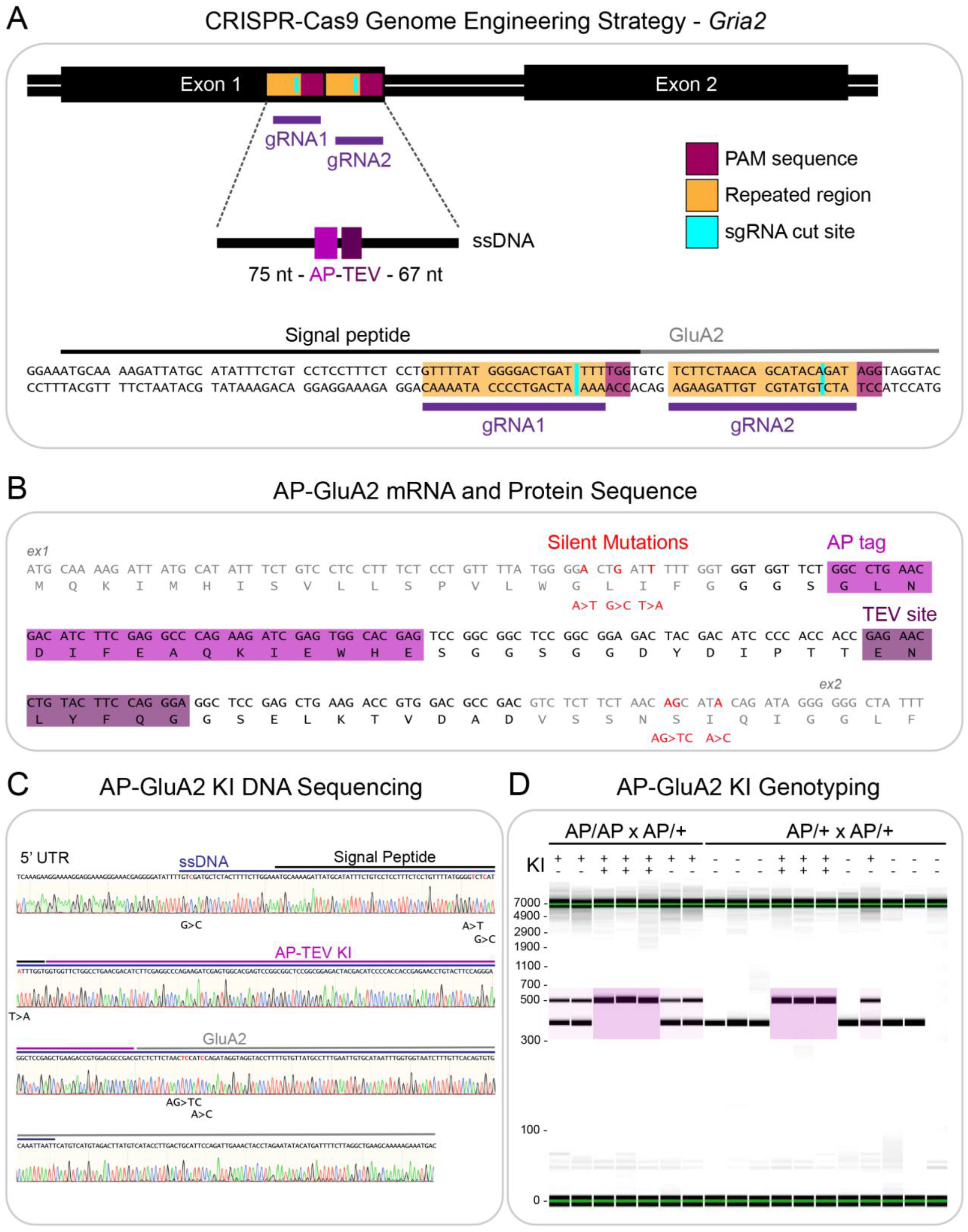
CRISPR-Cas9 strategy for AP-TEV knock-in on *Gria2*. (**A**). Overview of the genome editing approach to modify *Gria2* by CRISPR-Cas9 (Doudna and Charpentier, 2014), with a single stranded DNA donor template carrying the AP-TEV sequence targeted by two guide RNAs to PAM sequences in exon 1. (**B**). Modification of the N-terminal GluA2 mRNA/protein sequence by AP-TEV knock-in, including six silent mutations (three per gRNA target) encoded on the ssDNA template to avoid re-cutting after homology directed repair. Endogenous *Gria2* coding sequence is shown in grey, knock-in sequence is shown in black, and silent mutations are shown in red. The biotin acceptor peptide tag (AP) sequence is highlighted pink, and the tobacco etch virus (TEV) protease consensus sequence is highlighted purple. (**C**). Sanger sequencing of the male F0 founder. One unexpected silent mutation (G>C) was found in the 5’ UTR. (**D**). Example of PCR genotyping results obtained from homozygous or heterozygous KI and WT animals using capillary electrophoresis (LabChip GX). Sequencing primers (see Table S1) produce an amplicon of 494 bases from AP-TEV KI *Gria2*, highlighted in pink, and 350 bases from WT *Gria2*.

**Supplementary Figure S2:**
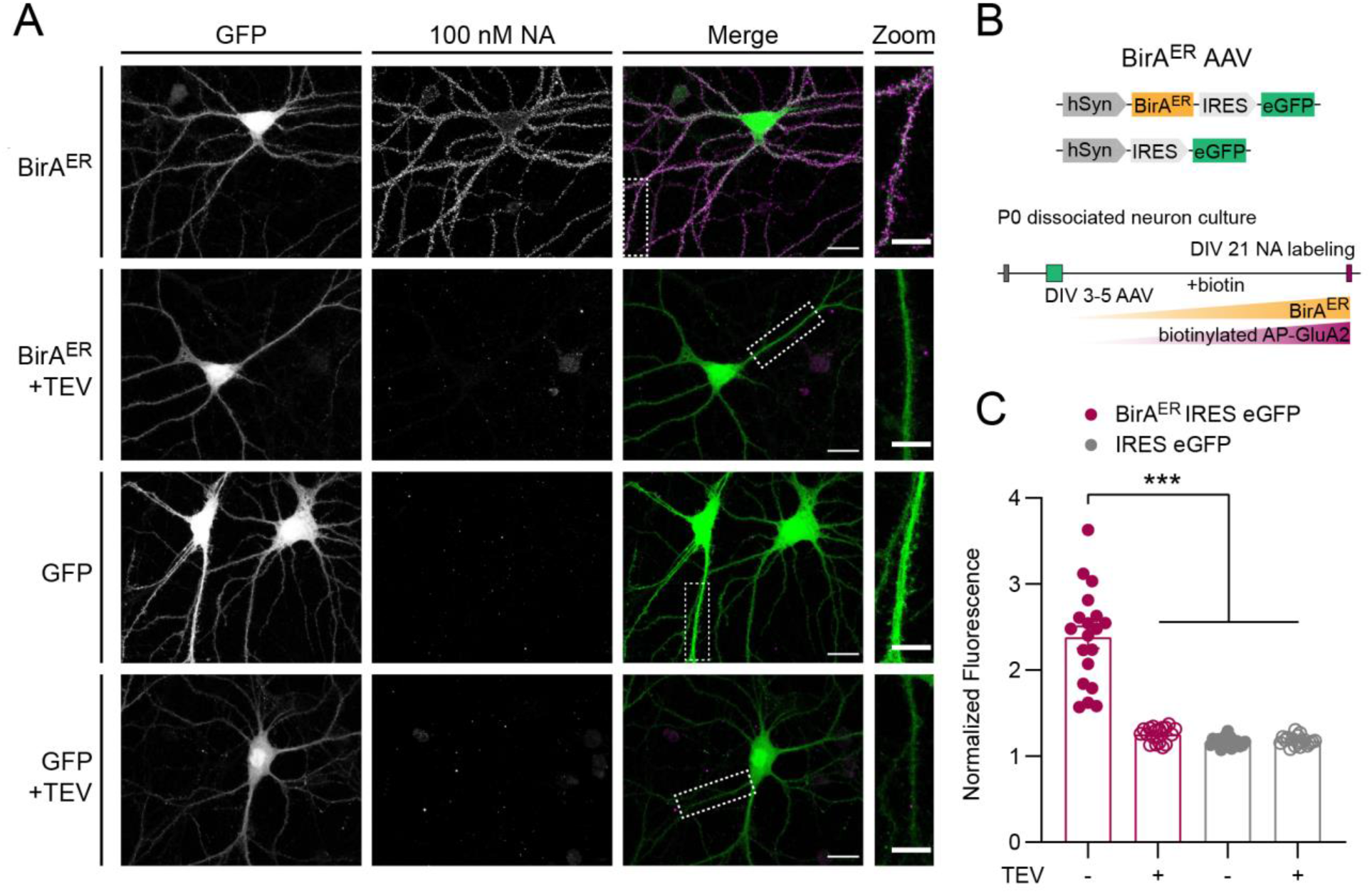
TEV protease cleaves the AP tag and reverses biotinylated AP-GluA2 surface labelling. (**A-B**). Primary hippocampal neuron cultures from AP-GluA2 KI mice transduced with AAV1 encoding BirA^ER^-eGFP or eGFP control and maintained for 21 days in media supplemented with 10 µM biotin. Cells were incubated with 100 nM NA conjugated to DyLight 633 to label surface localized bAP-GluA2, then incubated with a control buffer or 100 U TEV protease to cleave the AP tag and remove the NA surface label. (**A**). Representative confocal images of neurons transduced with BirA^ER^-eGFP or eGFP control AAV1, incubated with 100 nM NA, then with or without TEV. NA labelling is specific to AP-GluA2 KI with BirA^ER^ and without TEV. Scale bars, 20 and 10 µm. (**B**). Schematic representation of the experiment. (**C**). Normalized fluorescence intensity of NA-DyLight 633, coincident with the eGFP reporter. N≥17. \*\*\**P*≤0.0007 (Kruskal-Wallis test; F=47.18, *P*<0.0001; Dunn’s post-hoc test). Error Bars, SEM.

**Supplementary Figure S3:**
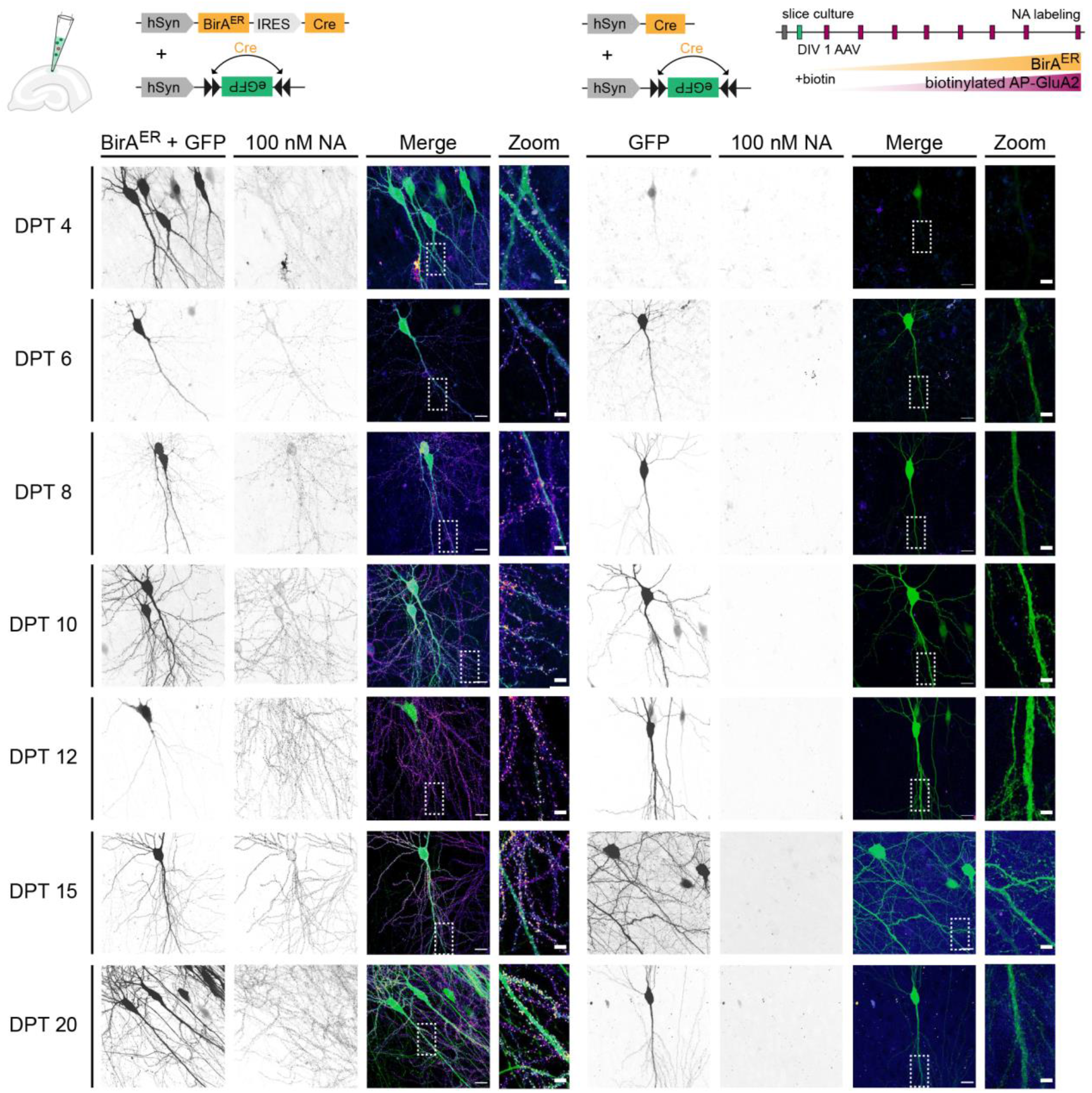
Time course of biotinylated AP-GluA2 expression with AAV-mediated expression of BirA^ER^. Organotypic slice cultures from AP-GluA2 KI mice transduced with AAV9 encoding BirA^ER^-Cre + FLEx eGFP or Cre + FLEx eGFP control in CA1 and maintained for 5-21 days in media supplemented with 10 µM biotin. Slices were incubated with 400 nM NA conjugated to DyLight 633 to label surface localized bAP-GluA2. Representative confocal images of CA1 pyramidal neurons in organotypic hippocampal slices transduced with BirA^ER^-Cre + FLEx eGFP (left) or Cre + FLEx eGFP control (right). DPT, days post transduction. Scale bars, 20 and 5 µm. Schematic representation of the experiment is shown above. N≥3. Normalized fluorescence intensity of NA-DyLight 633, coincident with the eGFP reporter, is shown in Figure 1E.

**Supplementary Figure S4:**
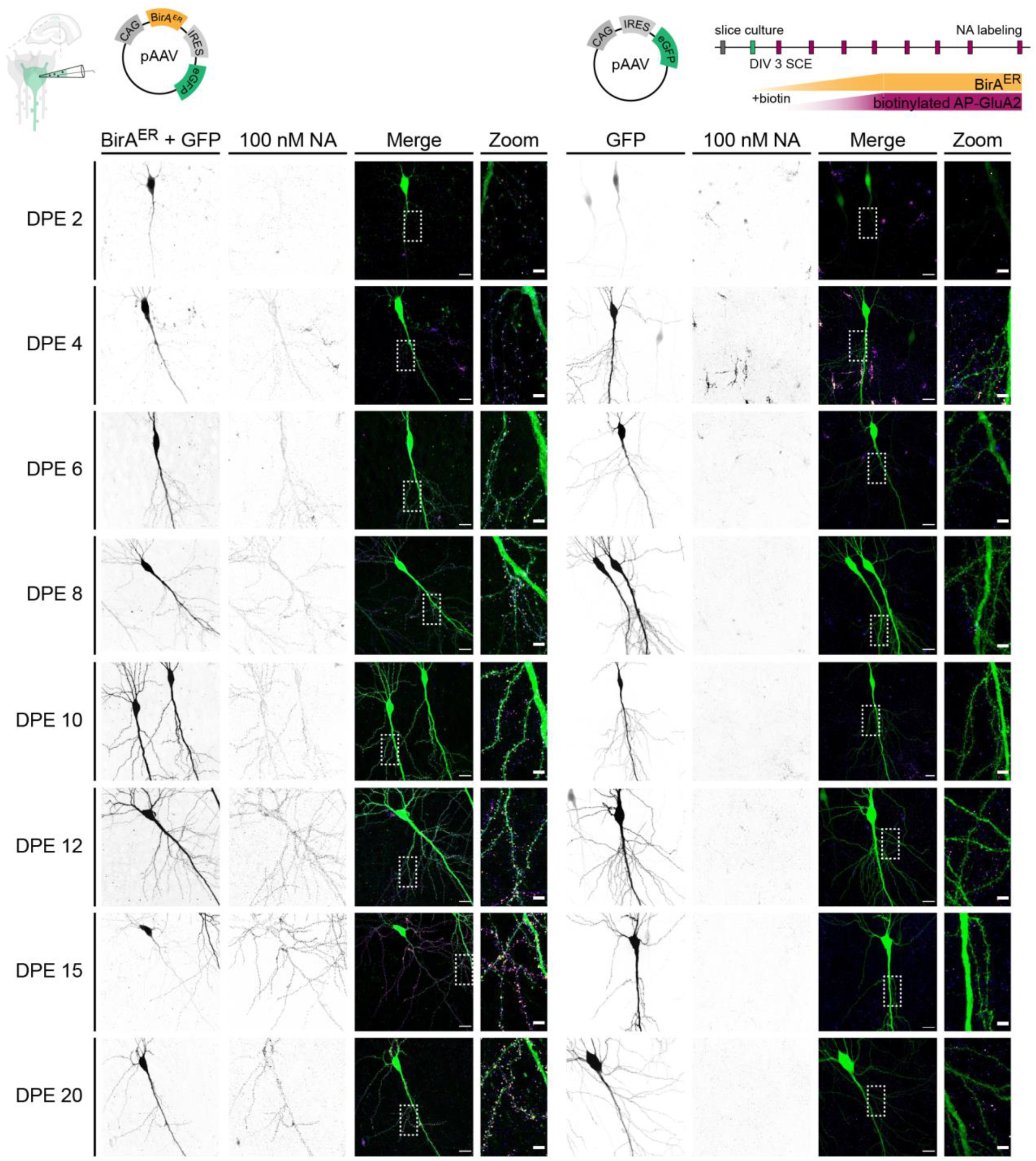
Time course of biotinylated AP-GluA2 expression with plasmid-mediated expression of BirA^ER^. CA1 pyramidal neurons in organotypic slice cultures from AP-GluA2 KI mice electroporated with 13 ng/uL pAAV plasmid encoding BirA^ER^-eGFP or eGFP control and maintained for 5-23 days in media supplemented with 10 µM biotin. Slices were incubated with 400 nM NA conjugated to DyLight 633 to label surface localized bAP-GluA2. Representative confocal images of CA1 pyramidal neurons in organotypic hippocampal slices electroporated with BirA^ER^-eGFP (left) or eGFP control (right). DPE, days post electroporation. Scale bars, 20 and 5 µm. Schematic representation of the experiment is shown above. N≥5. Normalized fluorescence intensity of NA-DyLight 633, coincident with the eGFP reporter, is shown in Figure 1G.

**Supplementary Figure S5:**
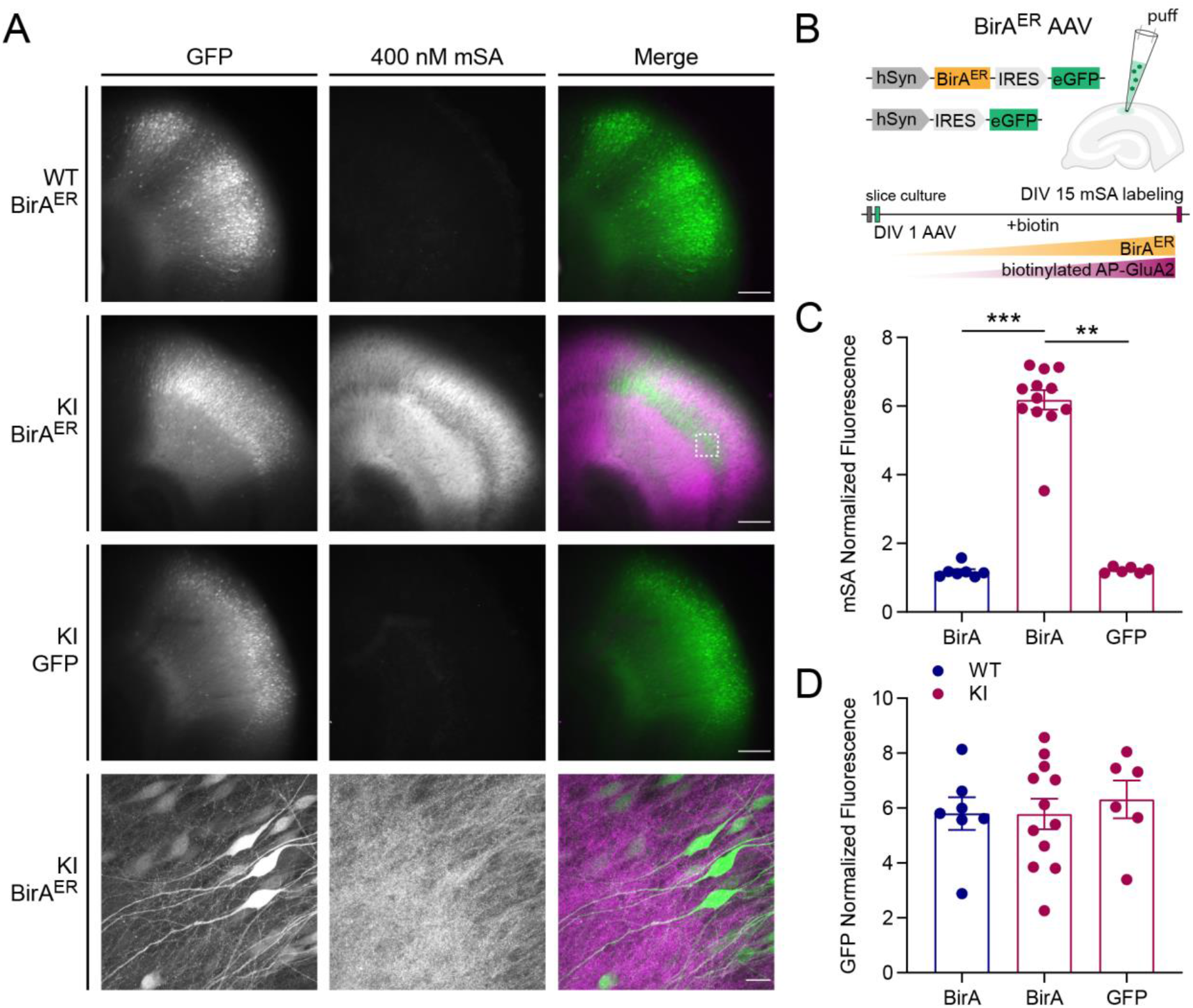
Specificity of mSA labelling of biotinylated AP-GluA2 in organotypic hippocampal slice cultures. (**A-B**). Organotypic slice cultures from AP-GluA2 KI or WT mice transduced with AAV9 encoding BirA^ER^-eGFP or eGFP control in CA1 and maintained for 15 days in media supplemented with 10 µM biotin. Slices were incubated with 400 nM mSA conjugated to Star 635P to label surface localized bAP-GluA2. (**A**). Representative widefield images of CA1 in organotypic hippocampal slices transduced with BirA^ER^ + eGFP or eGFP control AAV9 from WT or KI mice. mSA labelling is specific to AP-GluA2 KI with BirA^ER^. Bottom panel shows a representative confocal image from the boxed region. Scale bars, 200 and 20 µm. (**B**). Schematic representation of the experiment. (**C**). Normalized fluorescence intensity of mSA-Star 635P labelling in CA1, coincident with the eGFP reporter. N≥6. **-\*\*\**P*≤0.0072 (Kruskal-Wallis test; F=18.37, *P*=0.0001; Dunn’s post-hoc test). (**D**). Normalized fluorescence intensity of eGFP labelling in CA1. N≥6. *P*≥0.8237 (one-way ANOVA; F=0.1976, *P*=0.8222; Tukey’s post-hoc test). Error Bars, SEM.

**Supplementary Figure S6:**
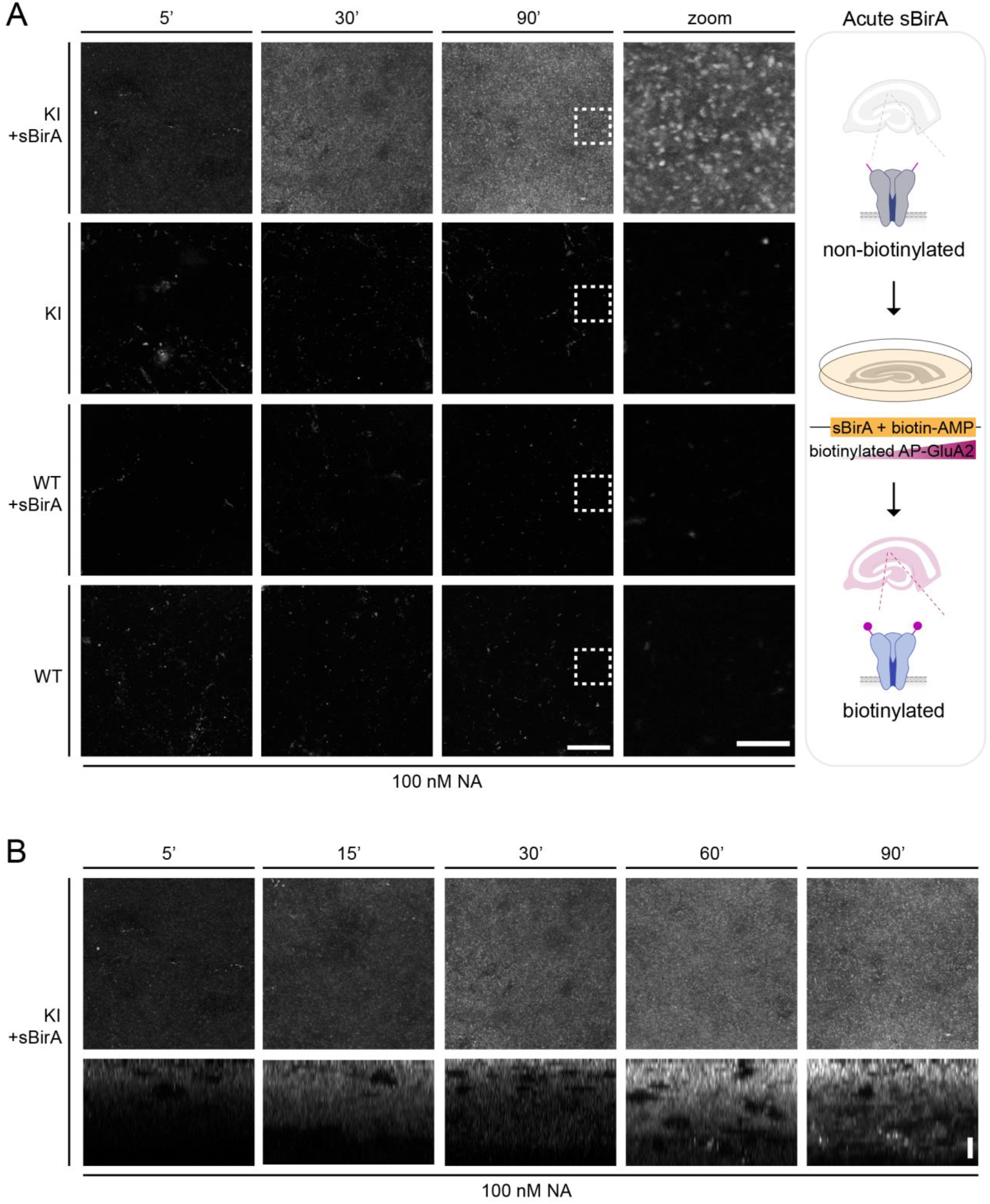
Time course of AP-GluA2 biotinylation with soluble BirA incubation. (**A**). Organotypic slice cultures from AP-GluA2 KI or WT mice incubated with 10 µM biotin-AMP and with or without 0.3 µM sBirA for 5, 30, or 90 min, then incubated with 400 nM NA conjugated to DyLight 633 to label surface localized bAP-GluA2. Representative confocal images from the stratum radiatum in organotypic hippocampal slices, as above. Scale bars, 20 and 5 µm. Schematic representation of the experiment is shown on the right. Normalized fluorescence intensity of NA-DyLight 633 is shown in Figure 1I. (**B**). Orthogonal projections (Y-Z) of NA-DyLight 633 labelled biotinylated AP-GluA2 after 5-90 min incubation with biotin-AMP and sBirA demonstrate progressive biotinylation of AP-GluA2 in depth over time. Scale bar, 10 µm.

**Supplementary Figure S7:**
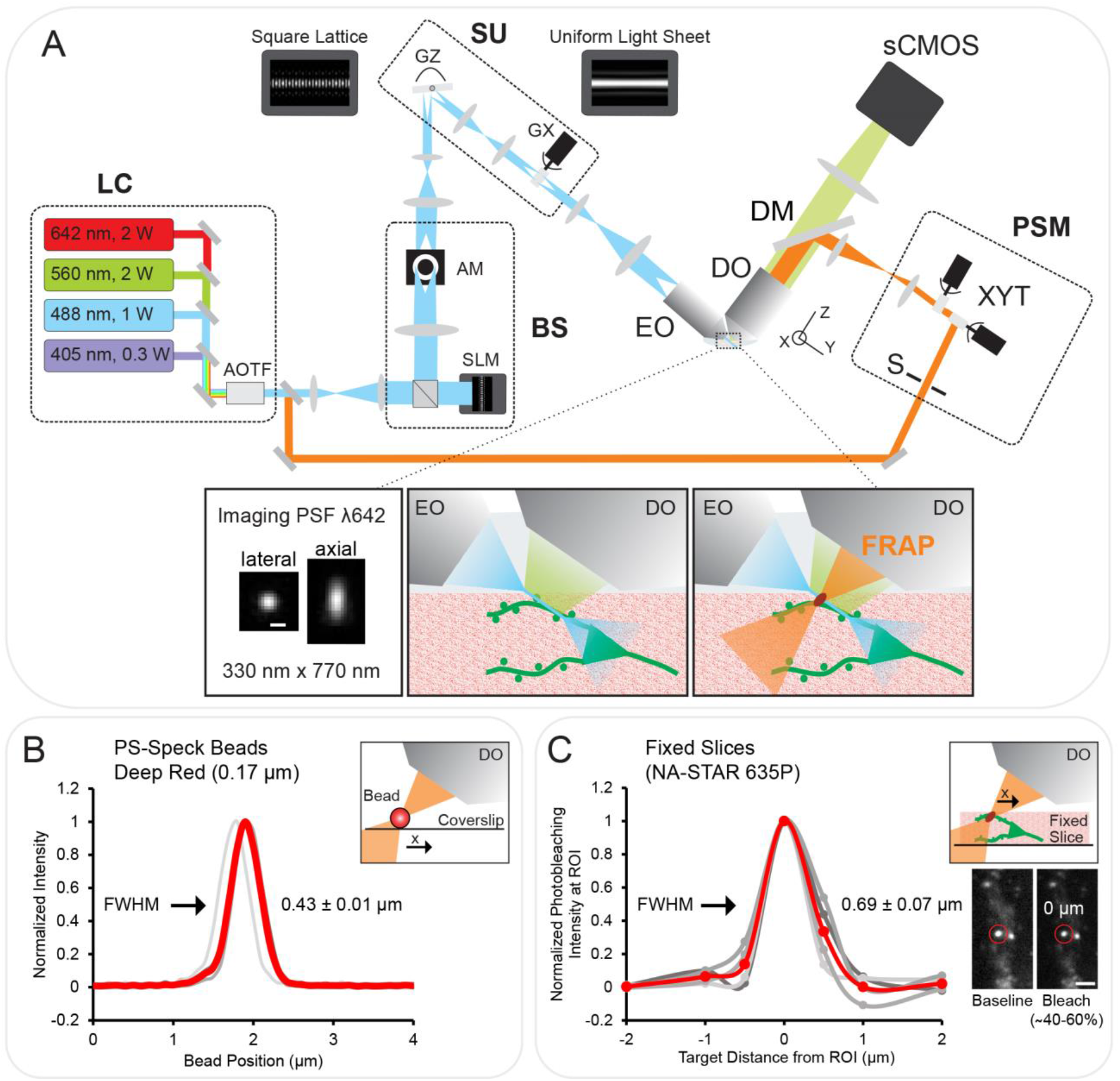
Photomanipulation with lattice light-sheet microscopy. (**A**). Lattice light-sheet microscope (LLSM) showing the original Betzig setup (Chen et al., 2014) with the added photomanipulation path. Excitation path is shown in blue, emission path in green, and PSM/FRAP path in orange. LC: four wavelengths laser combiner; AOTF: acousto-optic tunable filter; BS: beam shaper, composed of a spatial light modulator (SLM) and an annular mask (AM), generates the square lattice pattern; SU: scanning unit with Z and X galvos (GZ, GX), translates the square lattice pattern along the x and z axis, fast dithering with GX creates a uniform light sheet at the sample; EO: excitation objective; DO: detection objective; sCMOS camera detects fluorescence over the illuminated plane; PSM: photostimulation module, targets user-defined ROIs at high spatiotemporal resolution; S: shutter; XYT: galvo-based beam targeting system. Inserts show imaging point spread function at 642 nm excitation measured as 1/e^2^ radius (left), schematics of the sample with light sheet excitation and detected emission fluorescence (center), and the FRAP beam (right). The light sheet is a thin and extended plane (∼ 0.5 µm in z, >15 µm in Y, 100 µm in X) allowing fast 3D high resolution imaging. Scale bar, 0.5 µm. (**B**). FRAP dimension was measured by scanning a 0.17 µm diameter PS-Speck deep red bead (633/660 nm) across the FRAP beam focus (0.43 ± 0.01 µm full width at half maximum; FWHM). N=4, average profile shown in red. (**C**). FRAP dimension and efficiency in optically aberrating brain tissue was measured in fixed organotypic hippocampal slice cultures from AP-GluA2 KI mice that were transduced with BirA^ER^-Cre + FLEx eGFP AAV9 and incubated with 100 nM NA-Star 635P. The FRAP ROI was targeted at ± 0, 0.5, 1, and 2 µm from NA-labelled spines to measure the photobleaching profile in response to a ∼3 mW, 50 ms pulse at 642 nm (∼40-60% photobleach intensity; 0.69 ± 0.07 µm full width at half maximum; FWHM). N=4, average profile shown in red. Scale bar, 2 µm.

**Supplementary Figure S8:**
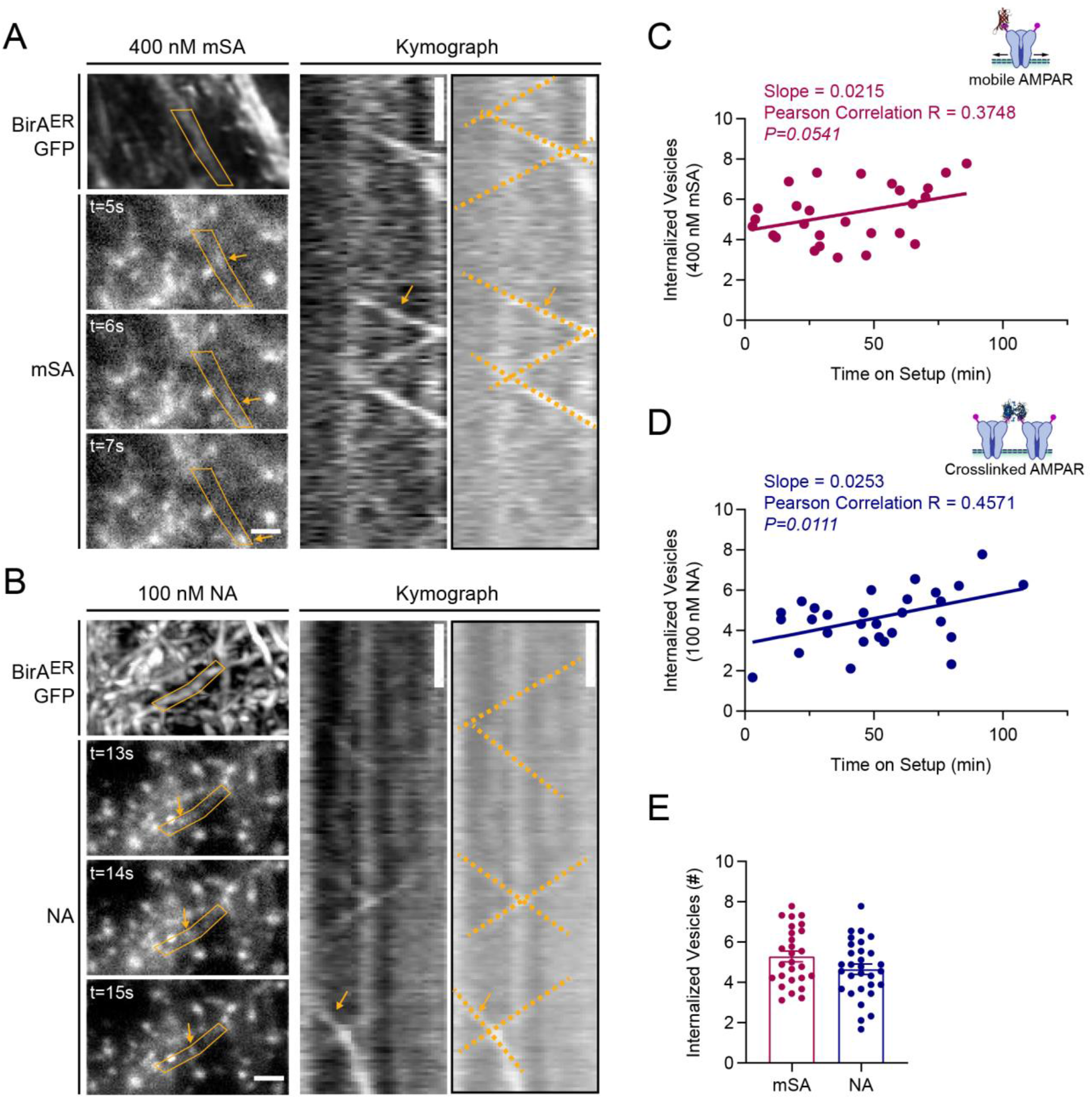
AP-GluA2 dendritic trafficking reveals extent of AMPAR internalization with monovalent or tetravalent biotin binding proteins. (**A-B**). Representative LLSM images of CA1 pyramidal neurons in organotypic hippocampal slice cultures from AP-GluA2 KI mice transduced with AAV9 encoding BirA^ER^ + eGFP in CA1 and maintained for 10-15 days in media supplemented with 10 µM biotin. Slices were incubated with 400 nM mSA-Star 635P (**A**) or 100 nM NA-Star 635P (**B**) to label surface localized biotinylated AP-GluA2. Live 3D volume stack of the eGFP reporter (top left) and continuous single plane acquisitions (10 Hz) of mSA or NA reveal intracellular trafficking events of AMPAR-containing internalized vesicles (arrows), quantified with kymographs along ∼5 µm of dendrite (right). Scale bars, 2 µm and 2 s. (**C-D**). Correlation of internalized vesicle counts vs time on setup for 400 nM mSA-Star 635P (**C**; Pearson R=0.3748, *P*=0.0541) or 100 nM NA-Star 635P (**D**; Pearson R=0.4571, \**P*=0.0111) labelled slices. (**E**). Mean number of internalized vesicles detected. N≥27. *P*=0.0944 (unpaired T-test). Error Bars, SEM.

**Supplementary Figure S9:**
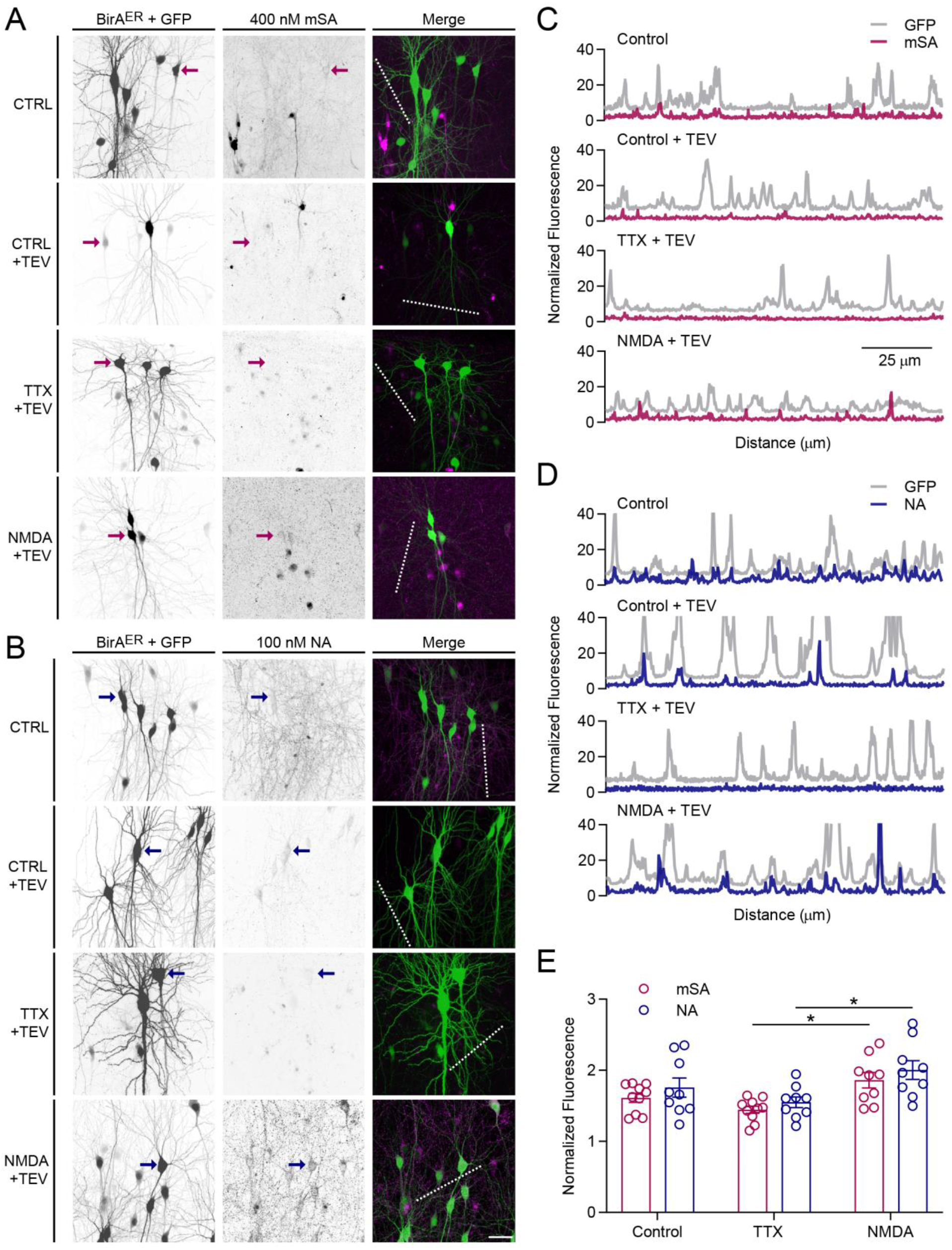
TEV proteolytic cleavage of surface AP-GluA2 reveals extent of AMPAR internalization with monovalent or tetravalent biotin binding proteins. (**A-B**). Representative confocal images of CA1 pyramidal neurons in organotypic slice cultures from AP-GluA2 KI mice transduced with AAV9 encoding BirA^ER^-Cre + FLEx eGFP in CA1 and maintained for 12-15 days in media supplemented with 10 µM biotin. Slices were incubated with 400 nM mSA-Star 635P (**A**) or 100 nM NA-DyLight 633 (**B**) to label surface localized bAP-GluA2, then incubated for 30 min in control conditions, with TTX (1 µM), or after NMDA treatment (30 µM, 3 min). Slices were then incubated with a control buffer or 100 U TEV protease (10 min) to cleave the AP tag and remove the mSA or NA surface label to reveal internalized AMPAR. Scale bar, 20 µm. (**C-D**). Linescans (dashed lines in A, B) reveal colocalization of mSA (**C**; Control -TEV: Pearson R=0.5356, \*\*\**P*<0.0001; Control +TEV: Pearson R=0.0998, \**P*=0.0223; TTX +TEV: Pearson R=0.0012, *P*=0.9789; NMDA +TEV: Pearson R=0.1083, \**P*=0.0132) or NA (**D**; Control -TEV: Pearson R=0.4490, \*\*\**P*<0.0001; Control +TEV: Pearson R=0.5265, \*\*\**P*<0.0001; TTX +TEV: Pearson R=0.0089, *P*=0.8394; NMDA +TEV: Pearson R=0.2807, \*\*\**P*<0.0001) with the eGFP reporter, and differential AMPAR internalization upon TTX or NMDA treatment (+TEV). (**E**). Normalized fluorescence intensity of mSA-Star 635P or NA-DyLight 633 after TEV proteolytic cleavage, coincident with the eGFP reporter. Crosslinking by tetravalent NA does not alter AMPAR surface distribution or recycling. N≥9. \**P*≤0.0400 (one-way ANOVA; F=4.5940, *P*=0.0016; Tukey’s post-hoc test). Error Bars, SEM.

**Supplementary Figure S10:**
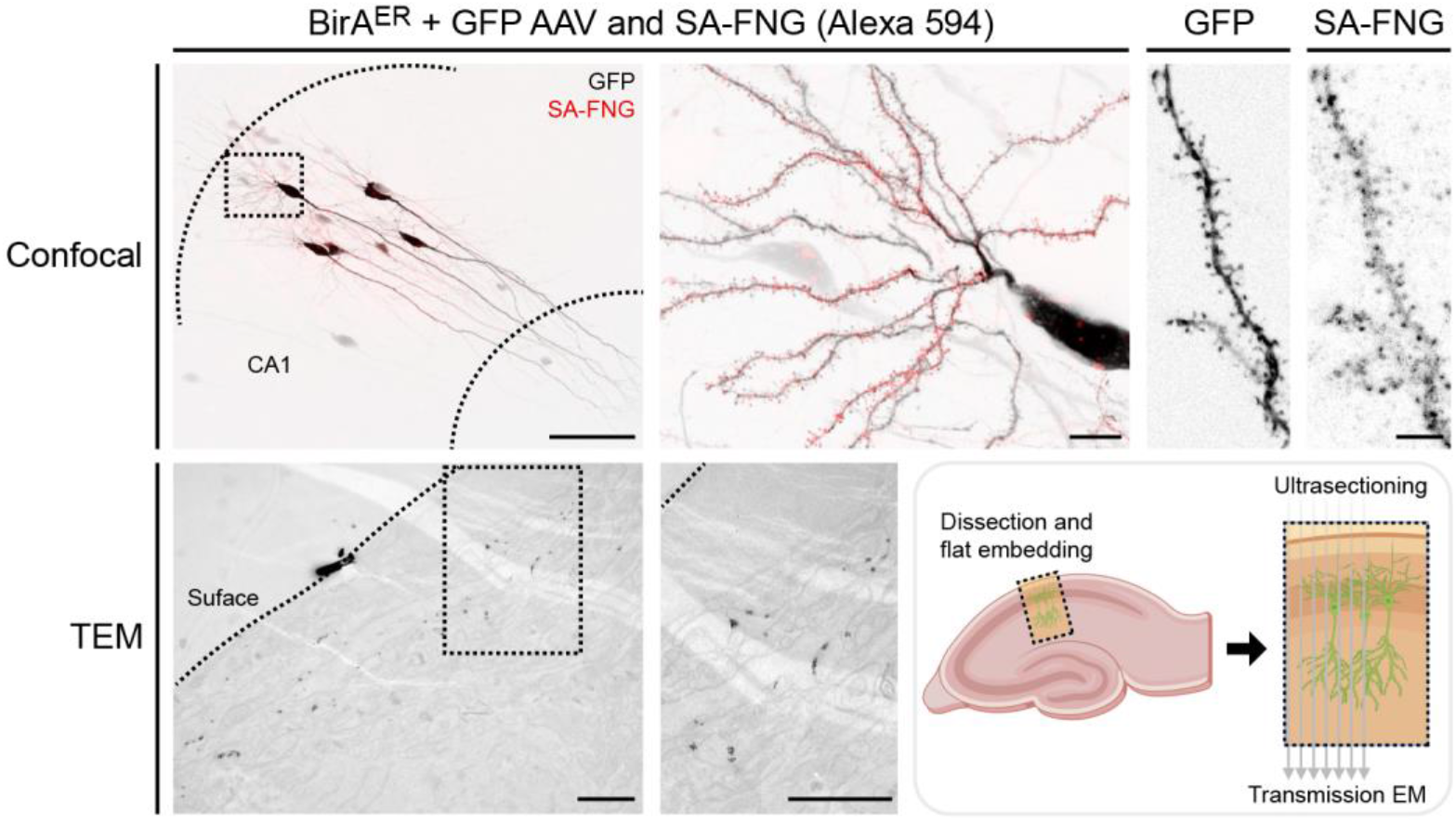
Sample preparation for TEM. Representative confocal images of organotypic hippocampal slice cultures from AP-GluA2 KI mice transduced with AAV9 encoding BirA^ER^-Cre + FLEx eGFP in CA1 and maintained for 15 days in media supplemented with 10 µM biotin, then incubated for 20 min with 100 nM StreptAvidin conjugated to AlexaFluor 594 FluoroNanoGold (SA-FNG) and fixed. Scale bars, 100, 10 and 5 µm (top panel). SA-FNG labeled slices were then processed and embedded for ultrasectioning of eGFP-positive ROIs. A low magnification image in TEM shows nanogold particle labeling after pre-embedding silver enhancement. Scale bars, 1 µm (bottom panel). Schematic representation of the experiment is shown on the bottom right. TEM synaptic micrographs are shown in Figure 3J, and quantification is shown in Figure 3K.

**Supplementary Figure S11:**
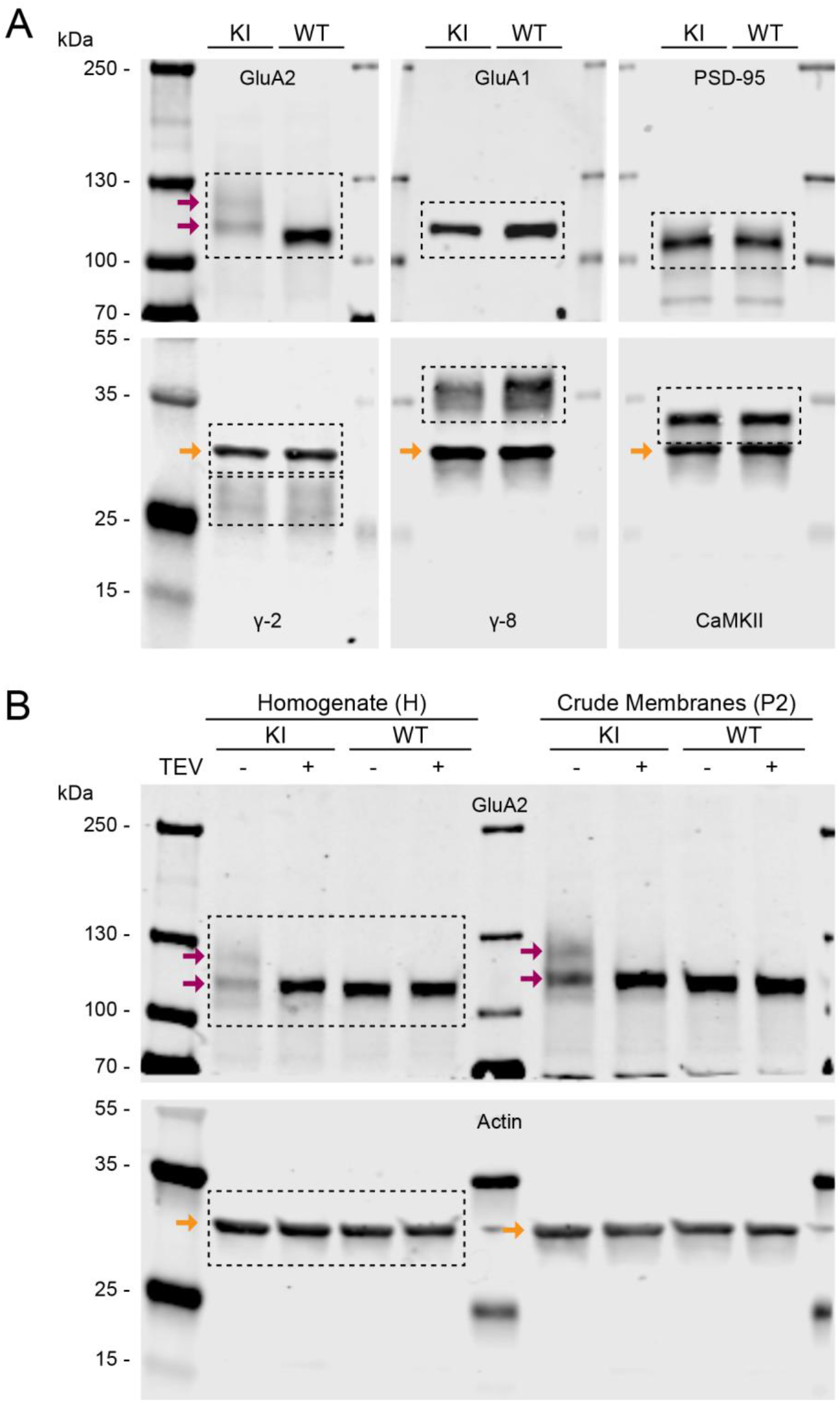
Characterization of AP-GluA2 and synaptic protein expression in KI and WT brain. (**A**). Representative Western blots of synaptic proteins in AP-GluA2 KI and WT whole brain protein samples. Double banding of GluA2 is observed for AP-GluA2 KI (red arrows). Boxed regions are shown in Figure 4A, and quantification is shown in Figure 4B. (**B**). Representative Western blot of the TEV proteolytic cleavage assay with whole brain lysate (H) or crude membrane fractions (P2) from KI and WT mice. Double banding of GluA2 is observed for AP-GluA2 KI without TEV (red arrows), and a single band with the same relative expression as WT is observed upon incubation with TEV protease. Boxed regions are shown in Figure 4C, and quantification is shown in Figure 4D. Membranes were cut between the 70 and 55 kDa markers to facilitate differential antibody labelling. β-actin was used as a loading control (orange arrows).

**Supplementary Figure S12:**
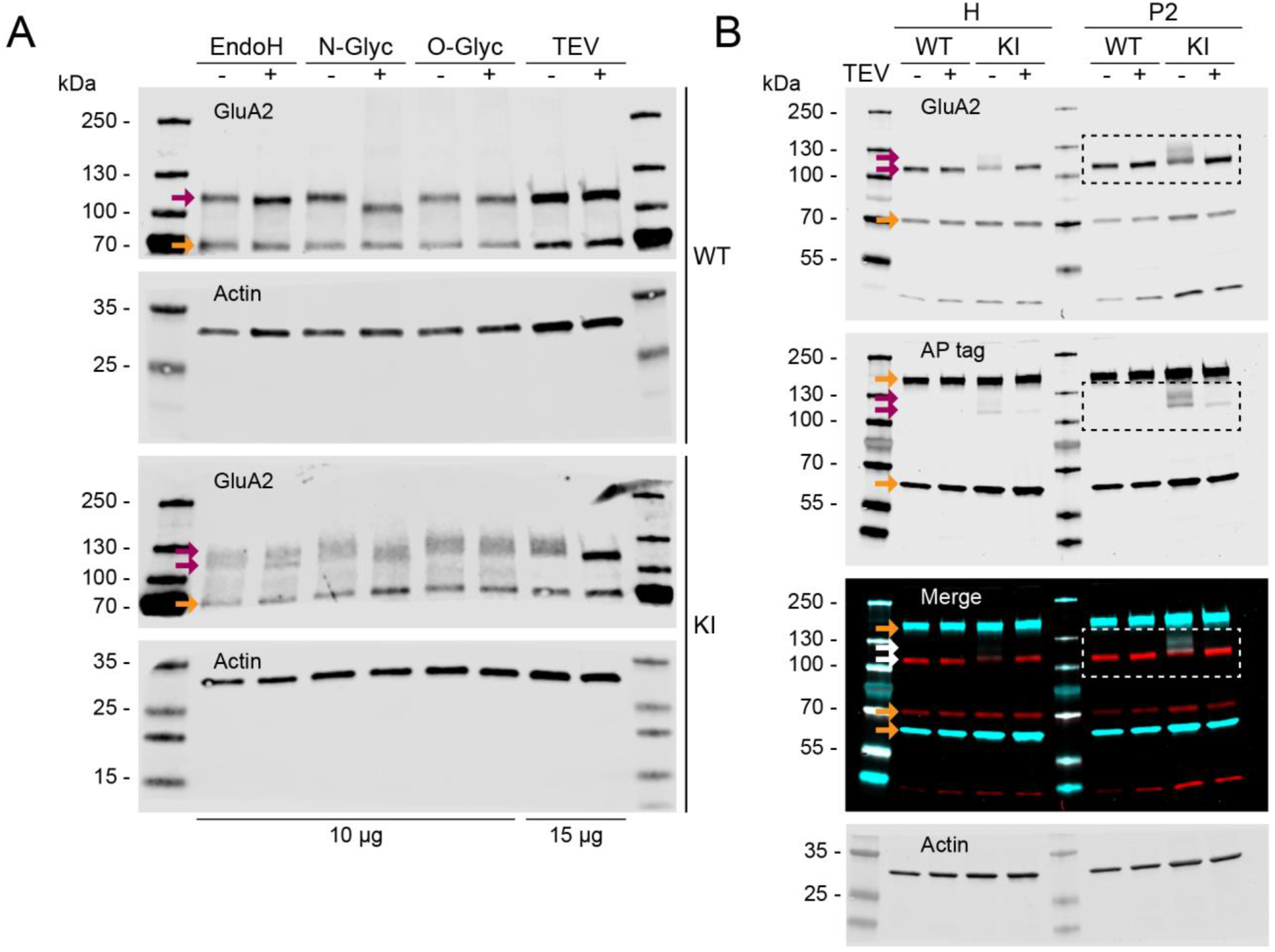
N-terminal degradation underlies the double banding pattern of AP-GluA2. (**A**). Representative Western blots of the GluA2 deglycosylation assay in WT and KI mice, protein samples were incubated with or without EndoH (N-glycan), PNGase F (N-Glyc; N-glycan), O-Glycosidase (O-Glyc; O-glycan), or TEV protease. Double banding of GluA2 is observed for AP-GluA2 KI (red arrows). Deglycosylation of GluA2 increases migration but does not affect double banding pattern of AP-GluA2. (**B**). Representative Western blots of AP tag and GluA2 (red arrows) in whole brain lysate (H) or crude membrane fractions (P2) from WT and KI mice, incubated with or without TEV protease. Boxed regions are shown in Figure 4E. Orange arrows indicate aspecific bands of the GluA2 or AP tag antibody. Membranes were cut between the 70 and 55 kDa markers to facilitate differential antibody labelling. β-actin was used as a loading control.

**Supplementary Figure S13:**
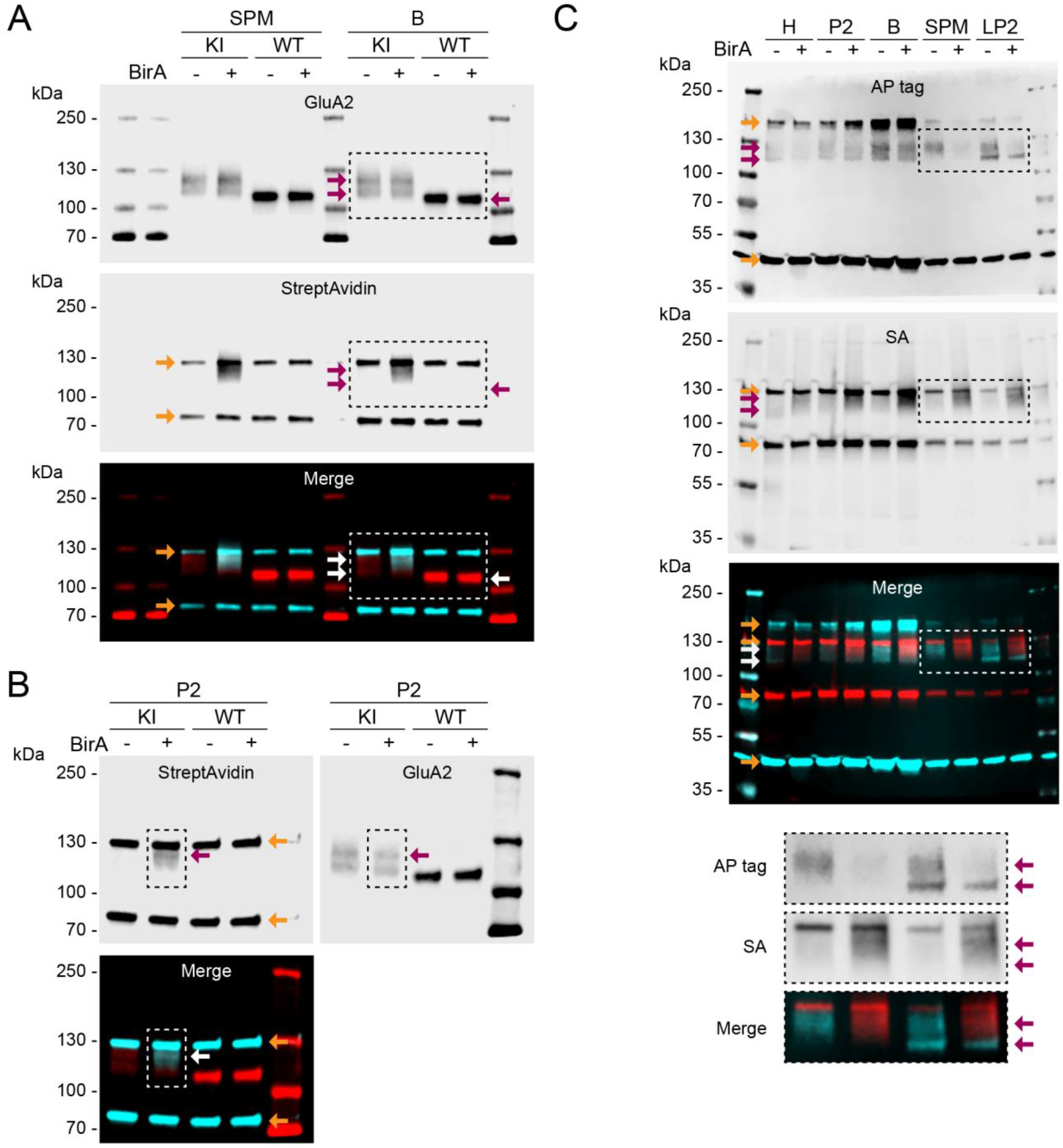
Biochemical characterization of AP-GluA2 in vitro biotinylation. (**A**). Representative Western blot of GluA2 in AP-GluA2 KI and WT synaptic plasma membrane (SPM) or synaptosome (B) fractions, incubated with or without soluble recombinant BirA. Double banding of GluA2 is observed for AP-GluA2 KI (red arrows). StreptAvidin binding of biotinylated AP-GluA2 is observed on the upper band. Orange arrows indicate StreptAvidin binding to endogenous biotin binding proteins pyruvate carboxylase (upper) and β-methyl crotonyl CoA carboxylase (lower). β-methyl crotonyl CoA carboxylase was used as a loading control. Boxed regions are shown in Figure 4G. Quantification of SA binding (+ relative to - sBirA), normalized to β-methyl crotonyl CoA carboxylase loading control, is shown in Figure 4K. (**B**). Representative Western blot of GluA2 and SA in crude membrane fractions (P2), as in (A), boxed regions used for linescan analysis are shown in Figure 4H. (**C**). Representative Western blot of AP tag in whole brain lysate (H), crude membrane fractions (P2), synaptosome (B), synaptic plasma membrane (SPM), or intracellular vesicle (LP2) fractions from AP-GluA2 KI mice, incubated with or without soluble recombinant BirA. Double banding of GluA2 is observed with the AP tag antibody (red arrows). StreptAvidin binding of biotinylated AP-GluA2 is observed on the upper band. Orange arrows indicate aspecific bands of the AP tag antibody or StreptAvidin binding to endogenous biotin binding proteins.

**Supplementary Figure S14:**
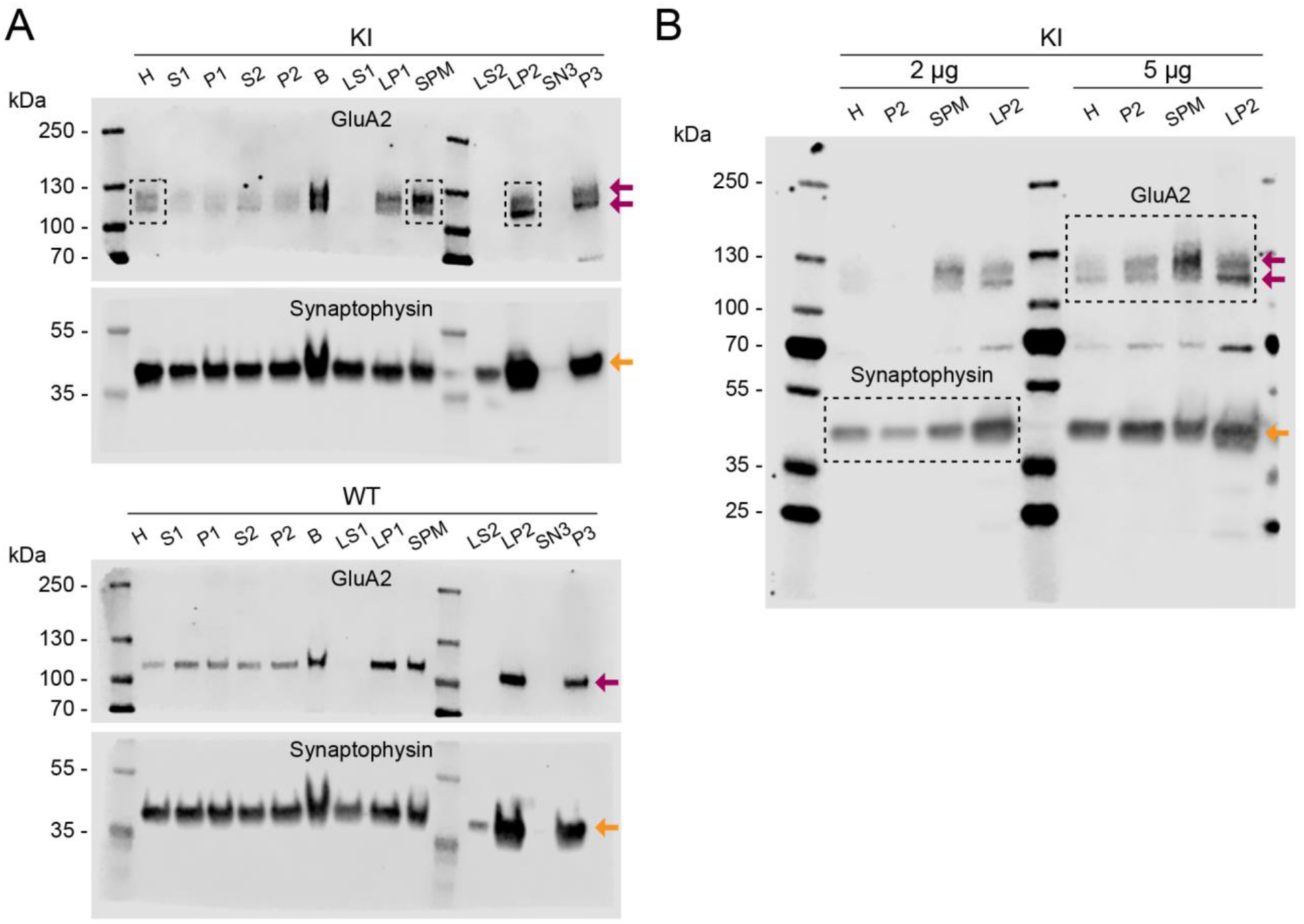
Subcellular fractionation reveals differential trafficking of AP-GluA2 protein isoforms. (**A**). Representative Western blots of GluA2 in AP-GluA2 KI and WT subcellular fractions (red arrows), as described in (De-Smedt-Peyrusse et al., 2018). Boxed regions used for linescan analysis are shown in Figure 4J, and quantification is shown in Figure 4L (H, homogenate; P2, crude synaptosomes, used to isolate SPM and LP2 fractions; SPM, synaptic plasma membrane; LP2; crude synaptic vesicles). Synaptophysin was used as a marker of intracellular vesicles (orange arrows), and is enriched in LP2. (**B**). Representative Western blot of GluA2 (red arrows) and Synaptophysin (orange arrow) in AP-GluA2 KI protein fractions of interest. Boxed regions are shown in Figure 4I. The upper GluA2 band (full length AP-GluA2, biotinylated) is enriched in the SPM fraction. The lower GluA2 band (AP-GluA2 degradation product, not biotinylated) is enriched in the LP2 fraction.

**Supplementary Figure S15:**
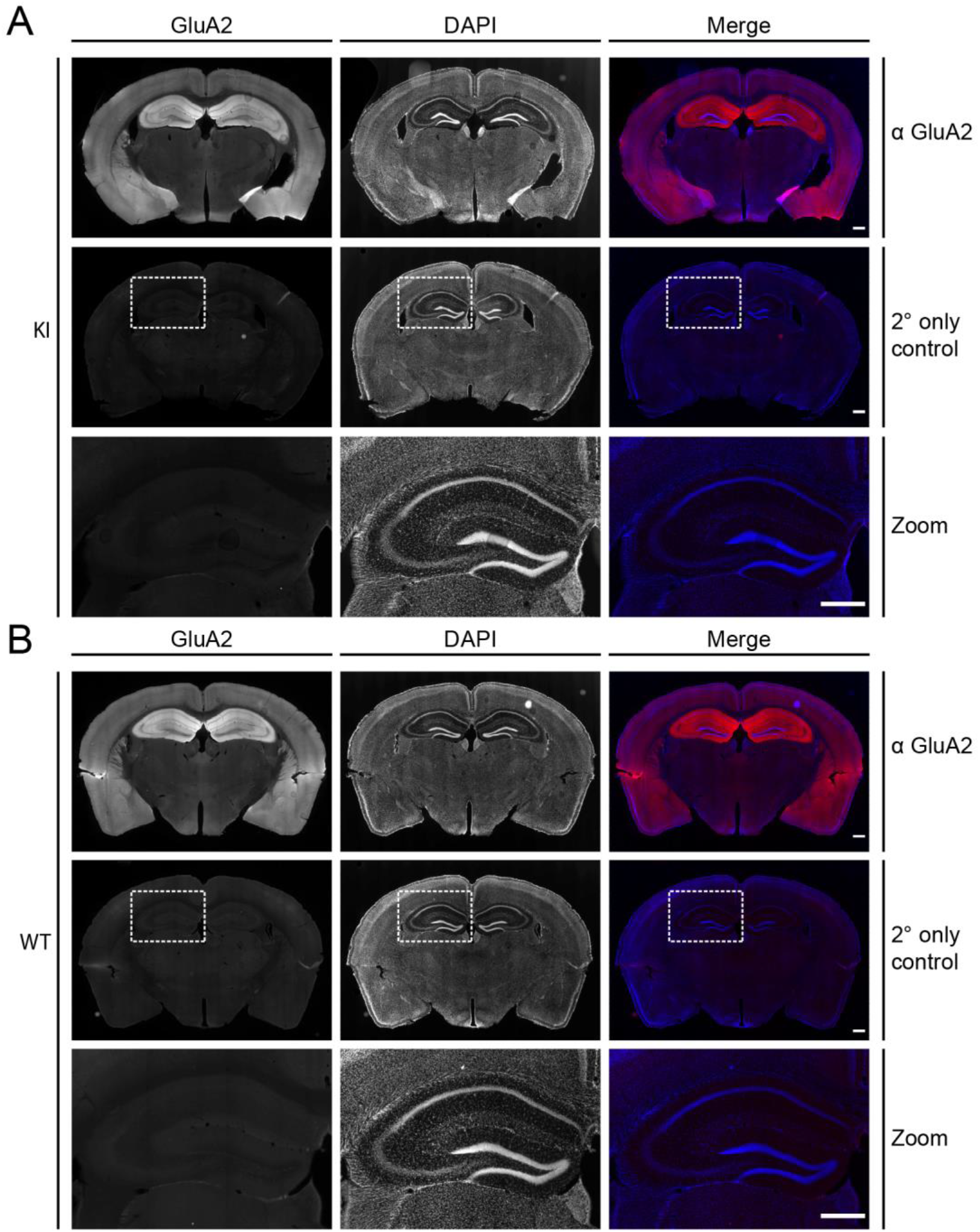
Secondary-only control for GluA2 IHC. (**A-B**). IHC characterization of GluA2 protein expression in frontal sections from AP-GluA2 KI (**A**) or WT mice (**B**), incubation without GluA2 primary antibody is shown to validate specificity of the fluorescent signal. Serial sections incubated with or without GluA2 primary antibody were used to normalize the fluorescence intensity signal in the data reported in Figure 3O-P. Scale bars, 500 µm.

**Supplementary Figure S16:**
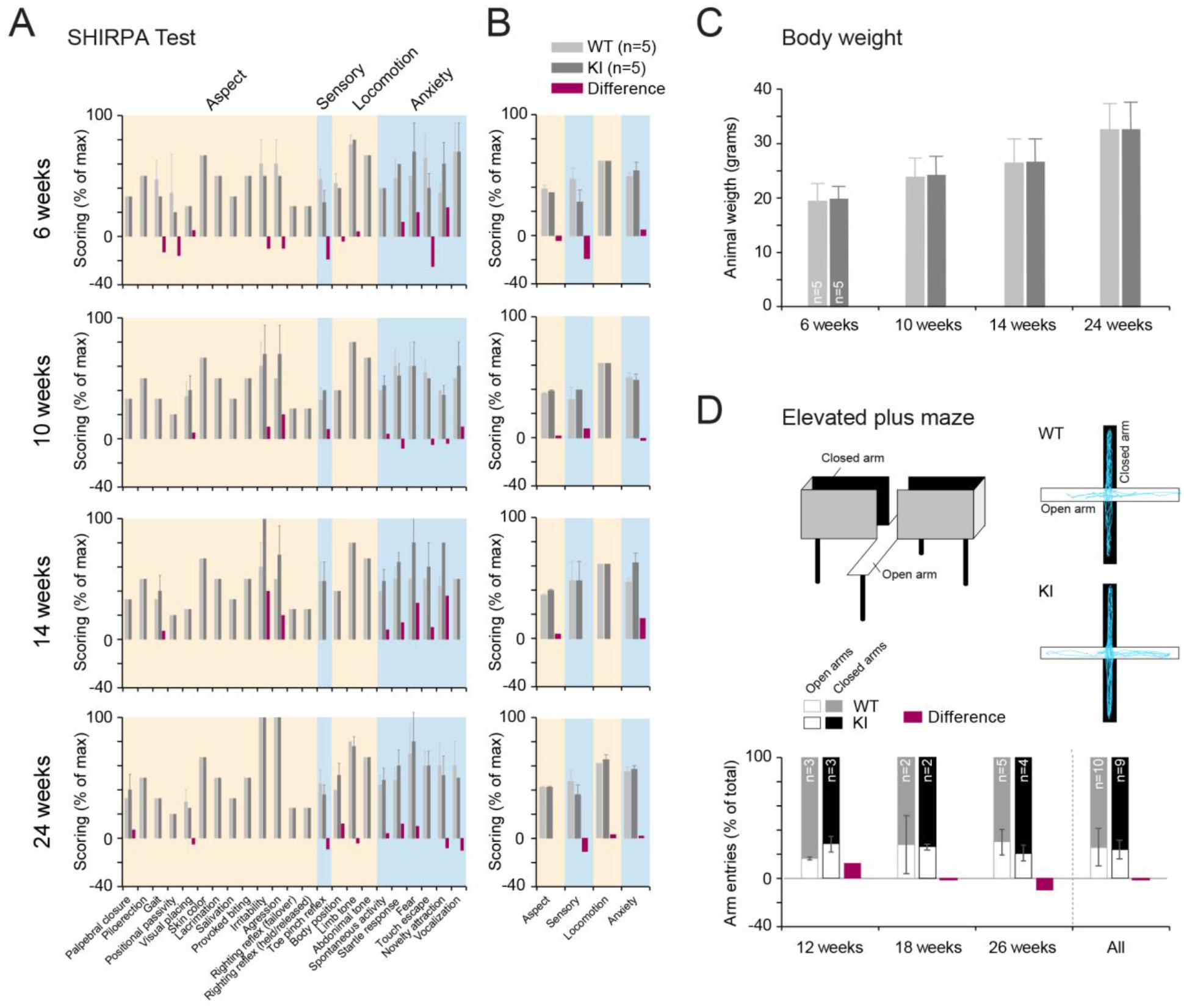
SHIRPA behavioral assessment of AP-GluA2 KI mice. (**A-B**). Behavioral assessment protocol for phenotype characterization of the AP-GluA2 KI mouse line. SHIRPA (SmithKline Beecham, Harwell, Imperial College, Royal London Hospital, phenotype assessment) longitudinal analysis of WT and KI mice at 4 time points (6, 10, 14 and 24 weeks). n=5. (**A**). Detailed screening behavior analysis for general aspect investigation, anxiety, locomotion and sensory abnormalities. Red bars represent the difference ratio between the two groups. (**B**). Grouped analysis for each aspect, Aspect-Sensory-Locomotion-Anxiety. Red bars represent the difference ratio between the two groups. (**C**). Body weight of WT and KI mice at 4 time points (6, 10, 14 and 24 weeks). *P*≥ 0.8781 (unpaired T-tests). (**D**). Schematic representation of the elevated plus maze test (top left). Deep lab cut tracking profile analysis of WT and KI mice in open and closed arms of the maze are shown in blue (top right). Percentage of open (white) and closed (shaded) arm entries observed in WT and KI mice at the 3 time points (12, 18 and 26 weeks). Red bars represent the difference ratio between the two groups. *P*=0.8246 (unpaired T-test; all data).

